# Elastomeric Pillar Cages Modulate Actomyosin Contractility of Epithelial Microtissues by Substrate Stiffness and Topography

**DOI:** 10.1101/2023.03.24.534106

**Authors:** Lisann Esser, Ronald Springer, Georg Dreissen, Lukas Lövenich, Jens Konrad, Nico Hampe, Rudolf Merkel, Bernd Hoffmann, Erik Noetzel

## Abstract

Cell contractility regulates epithelial tissue geometry development and homeostasis. The underlying mechanobiological regulation circuits are poorly understood and experimentally challenging. We developed an elastomeric pillar cage (EPC) array to quantify cell contractility as a mechanoresponse of epithelial microtissues to substrate stiffness and topography. The spatially confined EPC geometry consisted of 24 circularly arranged slender pillars (1.2 MPa, height: 50 μm, diameter: 10 μm, distance: 5 μm). These high-aspect-ratio pillars were confined at both ends by planar substrates with different stiffness (0.15 – 1.2 MPa). Analytical modeling and finite elements simulation retrieved cell forces from pillar displacements. For evaluation, highly contractile myofibroblasts and cardiomyocytes were assessed to demonstrate that the EPC device can resolve static and dynamic cellular force modes. Human breast (MCF10A) and skin (HaCaT) cells grew as adherence junction-stabilized 3D microtissues within the EPC geometry. Planar substrate areas triggered the spread of monolayered clusters with substrate stiffness-dependent actin stress fiber (SF)-formation and substantial single-cell actomyosin contractility (150 - 200 nN). Within same continuous microtissues, the pillar-ring topography induced bilayered cell tube growth. Here, low effective pillar stiffness overwrote local substrate stiffness sensing and induced SF-lacking roundish cell shapes with extremely low cortical actin tension (11 - 15 nN). This work introduced a versatile biophysical tool to explore mechanobiological regulation circuits driving low- and high-tensional states in developing and homeostatic microtissues. EPC arrays facilitate simultaneously analyzing the impact of planar substrate stiffness and topography on microtissue contractility hence microtissue geometry and function.

## Introduction

Epithelial tissue development and homeostasis are regulated by tightly-regulated forces that cells exert to and receive from their microenvironment [1]. These cellular forces modulate cell shape [2] and differentiation [3]. During development, distinct tensional force fields drive cell fate decisions [4], and branching morphogenesis of breast gland epithelia is characterized by local mechanical stress anisotropies [5]. Tissue deformation and folding arise from the contractility of individual cells in asymmetric tissue geometries where reciprocal feedback loops exist between tissue form and growth [6, 7]. Even within adult epithelial tissues, tension is adaptive to changing physiological demands. For instance, during the lifetime of breast epithelia, highly contractile alveoli evolve from a low contractile tissue state to produce and secrete milk and vanish again during involution [8].

Cell contractility is regulated by mechanical tissue cues, such as stiffness [9], topography and geometry [10, 11]. Cells sense and transduce these mechanical cues by integrin receptor-based cell-matrix adhesions. These focal adhesions (FAs) transmit forces via mechanical coupling to the actin cytoskeleton [12]. Myosin II coupling to actin stress fibers is essential for actomyosin-driven cell contractility [13]. Intracellularly, highly interconnected actomyosin networks transmit force over long distances [14, 15]. Intercellularly, force transmission is mediated by adherence junctions (AJs) coupled with the contractile actomyosin of neighboring cells [16]. Fundamental work demonstrated that local microenvironmental geometry modulates tissue development. Planar substrates micropatterned with a varying spacing of cell-matrix adhesion sites regulated cell growth and spreading [17]. More recently, epithelial clusters have been shown to adapt their cell size and alignment to differently shaped convex out-of-plane surfaces. This geometry-driven tissue shaping was functionally linked to changed actomyosin contractility [18]. Another work described the control of cell shape, actin SF formation and actomyosin tension by culturing cells within 3D microinches with different aspect ratios [19]. In addition to geometry, ECM stiffness is a fundamental regulator of normal and pathological tissue development [1]. Most of our knowledge about the impact of cellular stiffness sensing on cell contractility is based on traction force microscopy (TFM). Groundbreaking TFM techniques have been developed to measure forces that individual cells [20-22] and monolayered cell sheets exert on planar soft substrates of tunable stiffness [23, 24]. Such approaches revealed that cells join forces by transferring stresses through AJs during collective migration [25]. Advanced studies fabricated microneedles arrays with tunable spring constants to induce traction forces in single cells and monolayered clusters via sensing increased local substrate stiffness [26-28]. More recently, a three-dimensional (3D) TFM approach described that monolayered epithelial cell clusters exert 3D tractions on their microenvironment [29]. Moreover, hydrogel-based approaches reconstructed complex 3D traction fields of single cells [30, 31] and quantified the collective forces that growing tumor spheroids exert on their microenvironment [32].

Despite these sophisticated biophysical tools, quantitative TFM approaches are lacking to address how tension is maintained and regulated within defined 3D multilayered epithelial microtissues. To this end, we developed elastomeric pillar cages (EPCs) to quantify forces within engineered epithelial microtissues with areas of monolayered and bilayered shapes. Our device combines tissue engineering and laser-assisted nanosurgery with 3D cell force measurement in high-temporal and spatial resolution. We used highly resolving confocal microscopy to functionally link cytoskeletal reorganization with low tensional tissue states as a mechanoresponse to the EPC geometry. The present work introduced a versatile tool to study the modulation of microtissue contractility by substrate stiffness and topography.

## Material and Methods

### Preparation of microstructured casting molds

The EPC microstructures were written into lithography masks by electron beam lithography. Silicon wafers were preheated at 180 °C for 30 min and overlayed with a 45 μm thick layer of SU-8-25 photoresist (Microchem, Newton, MA)by spin coating at 1300 rpm. Subsequently, the photoresist layer was soft baked for 2 min at 65 °C followed by slow ramp heating to 90 °C (2 min) and an additional baking step (90 °C, 4 min). Ultraviolet photolithography (346 nm, source power 7 mW, 25 s) was used to transfer the EPC-photolithography mask into the photoresist layer. Wafers were post-baked for 7 min at 65 °C followed by slow ramp heating to 90 °C (5 min) and an additional baking step (90 °C, 3 min). The undeveloped photoresist was removed by washing with SU-8 developer (Microchem) for 8 min. Casting molds were hard-baked for 30 min at 180 °C.

### Preparation of elastomeric substrates

Elastomeric PDMS silicone rubber (Sylgard 184, Dow Corning, Midland, MI, USA) was used in mixing ratios of base oil and cross-linker oil 50:1 and 10:1 to generate elasticities of 15 kPa (pillar mold) and 1.2 MPa (bottom layer). Young’s modulus and Poisson’s ratio of elastomer samples were determined as described previously [33]. The casting mold was placed in a 3.5 mm cell culture dish, overlayed with 2 ml PDMS 10:1 w/w, treated with a mild vacuum, and cured at 60 °C for 16 h. Cured molds were peeled off, soaked in isopropanol (*p*.*a*) and cut into strips. Molds were treated with ultrasound bath (30 s in isopropanol). Pillar structures were dried without collapsing by using critical point drying and kept under a dry atmosphere until use. Planar bottom substrates (80 μm thickness, 15 kPa) were spin-coated on microscopic glass coverslips of 80 μm thickness (Menzel GmbH, Braunschweig, Germany), cured (16 h at 60 °C) and molds were gently placed on top. Coverslips with the assembled EPC strips were glued to the bottom of 3.5 cm μ-dishes (IBIDI, Gräfelfing, Germany) with predrilled 1.8 cm holes and cured (16 h at 60 °C). For visualization, elastomers were stained with the hydrophobic dye Vybrant DiD (V22887, 20 μmM in 70% ethanol, Thermo Fisher Scientific, Waltham, MA, US) at RT for 4 h. For cell force retrieval, 150 μL organic quantum dots solution (Qdot 655 ITK, Thermo Fisher Scientific) was mixed into 1 g PDMS (1:10) before placing it into the EPC casting mold.

### Cell maintenance

Non-transformed MCF10A cells were purchased from ATCC (Manassas, VA, USA) and maintained in culture dishes under standard culture conditions (37 °C, 5% CO_2_) in DMEM/F12 growth medium (Thermo Fisher Scientific) containing 5% horse serum (Thermo Fisher Scientific), 0.5 μg/mL hydrocortisone, 100 ng/mL cholera toxin, 20 ng/mL EGF, 10 μg/mL insulin (all Sigma Aldrich, St. Louis, MO, USA), 100 U/mL penicillin, and 100 μg/mL streptomycin (Thermo Fisher Scientific). MCF10A-rLVubi-LifeAct-TagRFP cells [34] were cultured as the MCF10Awt cells. MDA-MB-231 cells were grown in DMEM/F12 with 10% FCS (Thermo Fisher Scientific) and 100 U μL/mL penicillin and 100 μL/mL streptomycin. MCF-7 cells were cultured in RPMI1640 supplemented with 10% FCS, 1% sodium pyruvate (Thermo Fisher Scientific), 1% MEM non-essential AA (Thermo Fisher Scientific), insulin 10 μg μL/mL, and 100 U/mL penicillin and 100 μg/mL streptomycin. Keratinocyte cell line HaCaT was cultivated in DMEM high glucose (Sigma Aldrich) with 10% FBS, 100 U /mL penicillin, 100 μg/mL streptomycin and 1.8 mM calcium. Two days before experiments calcium concentration of 2.5 mM was supplemented to induce cell-cell-contact formation. Cardiomyocytes and myofibroblasts were freshly isolated from embryonic rats as previously described [35]. For differentiation to myofibroblasts, cells were cultivated one week before use. Both cell types were cultured in F10 Ham Media supplemented with 10% FBS, 100 U /mL penicillin and 100 μg/mL streptomycin, Insulin (1 mg/ml), transferrin (0.55 mg/ml) and sodium selenite (0.5 μg/ml) (Sigma Aldrich).

### Cell seeding in EPC arrays

EPC surfaces were functionalized with fibronectin (FN) (20 μg/mL in PBS). For precise seeding of cells into the pillar rings, borosilicate microcapillaries (outside diameter: 1.0 mm, inside diameter: 0.72 mm, Hilgenberg, Malsfeld, Germany) were used. The front end of the capillary was forged to an outer diameter of 15 μm using a capillary puller (Sutter Instrument, Novato, USA) and bent to an angle of 45° using a homemade microforge. Cells were detached from cell culture flasks with trypsin/ EDTA (0.05%), resuspended in growth media and loaded into the capillary (5×10^5^ cells / ml). Cell injection was performed by using a micromanipulator (Inject Man) and an oil-driven microinjector (CellTram vario) (both Eppendorf, Wesseling, Germany).

### Immunofluorescence staining

Cells were washed with cytoskeleton buffer (CB: 5 mM EGTA, 5 mM glucose, 10 mM MES, 5 mM MgCl_2_, 150 mM NaCl, 1 g/L streptomycin; Sigma Aldrich) at RT and fixed with 4% paraformaldehyde (Sigma Aldrich) in CB for 20 min (RT). After quenching with 100 mM glycine in CB for 30 min, samples were permeabilized with 1% Trition X-100 (Sigma Aldrich) for 20 min at RT. Non-specific antibody binding was blocked for 2 hours at RT with CB containing 5% milk powder, 0.1% BSA, 0.2% Triton X-100, and 0.05% Tween 20 (Sigma Aldrich). Samples were incubated overnight at 4 °C with primary antibodies anti-E-cadherin (mouse clone 36 BD610182, BD, Heidelberg, Germany), anti-paxillin (mouse clone 5H11 AHO0492, Thermo Fisher Scientific), anti-phospho-myosin light chain II 3671 (Cell Signaling, Danvers, MA, USA), in 1% blocking buffer (CB). Secondary antibodies conjugated with fluorescent dyes Alexa Fluor 405, Alexa Fluor 488, Alexa Fluor 546 (Thermo Fisher Scientific) and Atto 633 (Sigma Aldrich) were diluted in 1% blocking buffer (CB) and applied to the sample for 60 min at RT in darkness. Phalloidin-Alexa Fluor 488 (Thermo Fisher Scientific) staining was performed in the same step. Nuclei were counterstained with Hoechst 33342 (Thermo Fisher Scientific) for 60 min at RT. Washing steps were performed with CB. Samples were visualized and imaged using a Zeiss C-Apochromat water immersion objective lens (40x, NA = 1.4) on a confocal laser scanning microscope 880 (cLSM880) with ZEN 2.3 software (Carl Zeiss, Jena, Germany).

### Scanning electron microscopy

Scanning electron microscopy was performed using a Zeiss Leo 1550 scanning electron microscope (Carl Zeiss, Jena, Germany). The EPC array molds were coated with a 2 – 4 nm iridium layer using a sputter coater (Cressington 208 HR/MTM20, Watford, UK). Scanning electron microscopy was performed at 10 kV using the SE2 or InLense detector.

### Imaging of living and fixed cells

LCI was performed at 37 °C and 5% CO_2_ (cell incubator XL, Zeiss, Germany) with an inverse confocal laser scanning microscope (LSM880 with Airyscan detector) that used C-Apochromat water immersion autocorr objective lens (40x, NA = 1.2) (Carl Zeiss, Jena, Germany). Living cardiomyocytes, myofibroblasts and HaCaT cells were labeled with 100 nM MitoTracker Red FM (Thermo Fisher Scientific).

### Cell ablation

Cells were ablated using the laser ablation system UGA-40 Firefly with a 355 nm laser with 42 mW output power (Rapp Optoelectronic, Wedel, Germany) equipped to a cLSM880. For single-cell ablation, laser power of 2.5% (30 s) and for microtissue ablation of 3.5% (1.47 mW for 45 s) was used.

### 2D Traction force microscopy and EPC force retrieval

2D traction forces were analyzed on 15 kPa TFM substrates coated with FN (20 μg/mL in PBS). Substrate deformations were visualized by tracking 0.2 μm fluorescent beads (FluoroSpheres carboxylated crimson beads, Thermo Fischer) as described [36]. Marker bead recording started 5 min pre-cut and ended 20 min post-cut. Maps of cell-induced traction stresses were calculated by regularized least square fitting to the mechanical response of an elastic layer of 80 μm thickness on rigid substrates [21, 37]. For cell force calculation of pillar-bound cell clusters, image recording started 2 min pre-cut until 25 min post-cut (image interval: 30 s). The imaging focus plane was set 5 μm above the bottom substrate layer. For myofibroblasts, image recording started 1 min pre-cut and ended 3 min post-cut. For cardiomyocytes, pillar displacements were recorded for 120 s with an image interval of 90 ms. An in-house developed Python (version 3.7) program was used to track pillar displacements based on the Qdots pattern using cross-correlation as previously described [21]. Pillar positions at t = 0 s served as a reference to calculate displacement vectors for all time points. Cell forces were calculated from pillar displacements using the analytical approximation, based on previous work [21, 38-43] and as described in detail in Appendix A1.

## Statistical analyses

All measured values were plotted. The two-tailed nonparametric Kolmogorov-Smirnov-Test was performed for statistical data analyses of 2D traction forces of HaCaT and MCF10A cells. All tests were performed using GraphPad Prism version 9.5.1 (GraphPad Software, La Jolla, CA). The two-tailed nonparametric Kruskal-Wallis-Test was applied for significance tests of measured cell forces. Mean values are plotted with a 95% confidence interval (95% CI). The p-values were defined as follows: n.s.: p ≥ 0.05; *: p < 0.05; **: p < 0.01; ***: p < 0.001; ****: p < 0.0001).

## Results

### Microfabrication of EPCs for 3D epithelial cell cultures

We fabricated elastomeric pillar cages (EPC)s to analyze multicellular clusters in a mechanically defined 3D environment and to steer the 3D growth of multicellular cell assemblies. Micro-structured wafers containing two arrays of 40 EPCs were fabricated by soft replica molding (Figure 1a). These ready-to-use samples allowed for long-term cell cultivation and multiposition live cell imaging (LCI) experiments in high spatial and temporal resolution (Figure 1b). The upside-down view details the EPC mold (1.2 MPa) before assembling with the bottom substrate layer. An EPC consists of 24 pillars interspaced by stabilization bars to enable multiposition experiments. A large diffusion channel ensured optimal passive diffusion of growth media during long-time cell culture experiments (Figure 1c). Figure 1d highlights the narrow arrangement of high-aspect-ratio (AR = 1:5) pillars. This high AR led to occasional pillar-pillar adhesion at the soft bottom layer. To create spatially confined cage-like structures, the pillar rings and the planar top and bottom layers seamlessly connected (Figure 1e). Cell adhesion throughout the topography was realized by homogenous surface functionalization with the ECM protein fibronectin (FN) (Figure 1f). Self-forged glass capillaries were used for controlled cell injection through the pillar gaps (5 μm) into the lumen (Figure 1g, and Movie S1).

**Figure 1.**
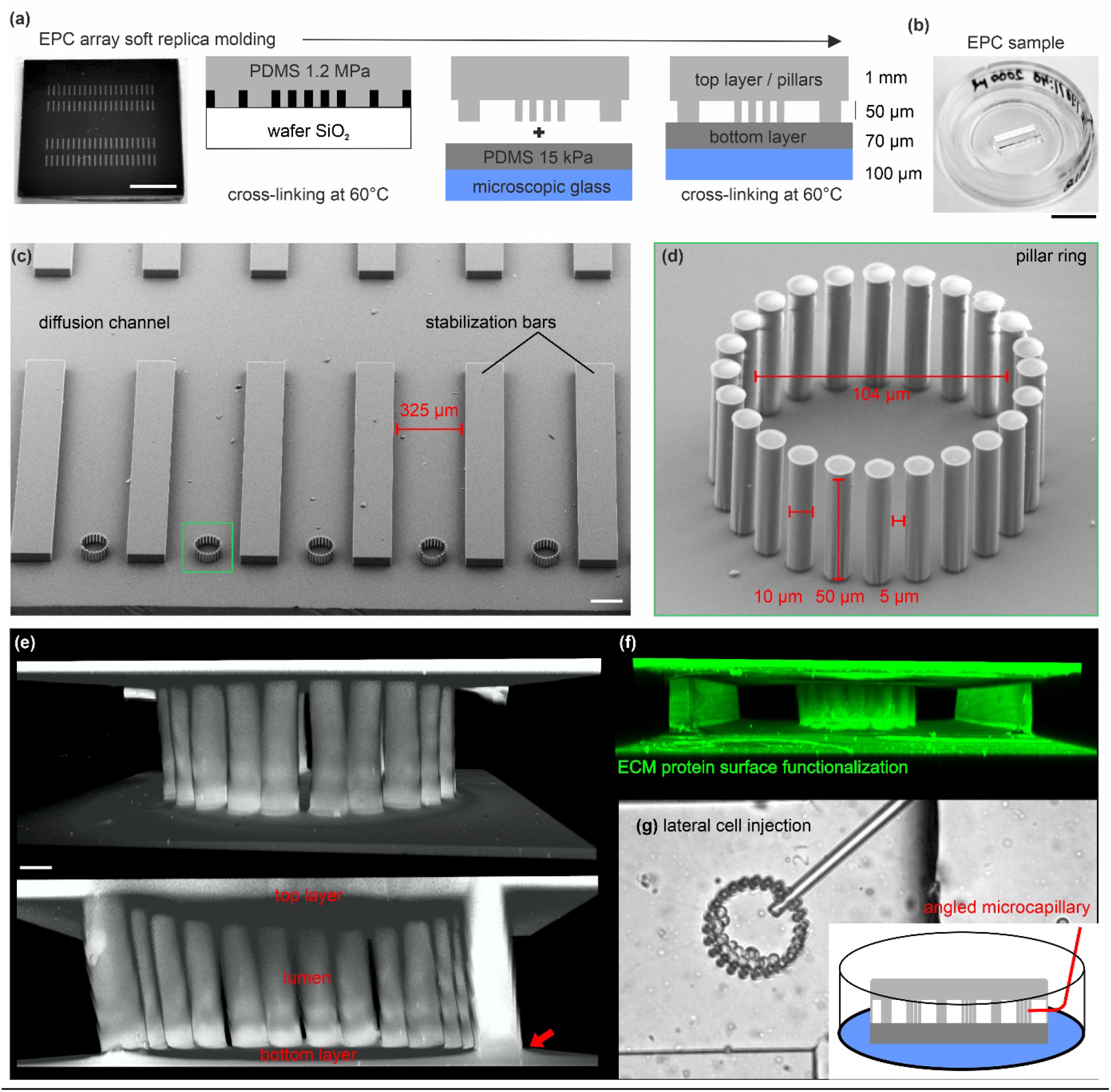
Microfabrication of EPC arrays for 3D epithelial cell cultures: (a) Outline for the soft replica molding process to produce EPC arrays. A silicon wafer microstructured with SU-8 resist was used as mold cast for the pillar ring structures. The fabrication process is described in detail in the materials and methods section. Scale bar = 0.5 cm. (**b**) The PDMS silicone rubber topography is mounted in a microscopic glass slide for live cell imaging. A rubber strip with two rows of 20 arrayed pillar rings is used per dish. Scale bar = 1 cm. (**c**) Scanning electron micrograph shows the array topography (upside down orientation). (**d**) gives a detailed view of a single ring structure with 24 high-aspect ratio pillars. Scale bar = 100 μm. The zoom-in micrograph highlights the high aspect ratio pillars. (**e**) 3D reconstructed image stack of a fully assembled ring structure confined by the planar bottom and top layers. The red arrow indicates the seamless material connection. The elastomer was stained with a hydrophilic fluorescent dye (DiD) for visualization. Scale bar = 10 μm. (**f**) A 3D reconstructed image stack of an EPC demonstrates the homogeneous ECM-surface functionalization with TRITC-labeled FN protein (20 μg / mL). (**g**) shows lateral cell injection into a pillar ring using self-forged glass microcapillaries.

### The EPC geometry induces the growth of mono-and bilayered epithelial microtissues

We aimed to grow 3D epithelial microtissues within our confined pillar scaffold. For this purpose, human keratinocytes (HaCaT) originated from adult skin [44], and non-transformed MCF10A cells derived from benign mammary breast gland tissue [45] were laterally injected into the lumen of the pillar cages and cultured for seven days. Four hours post-seeding, cells adhered equally to the planar bottom surface and pillars. After three days, circular cell-cluster localized along the pillar rings (Figure 2a and b). Both cell types formed double-layered tissue tubes after seven days. Here, the cells at the inner and outer pillar sides contacted each other through the 5 μm pillar gaps (Figure 2a, red arrow). The shown representative MCF10A cell tube consisted of 80 cells with direct pillar contact. Figure 2c demonstrate the growth of confluent monolayered cell clusters on the planar top and bottom substrates (Figure 2c).

**Figure 2.**
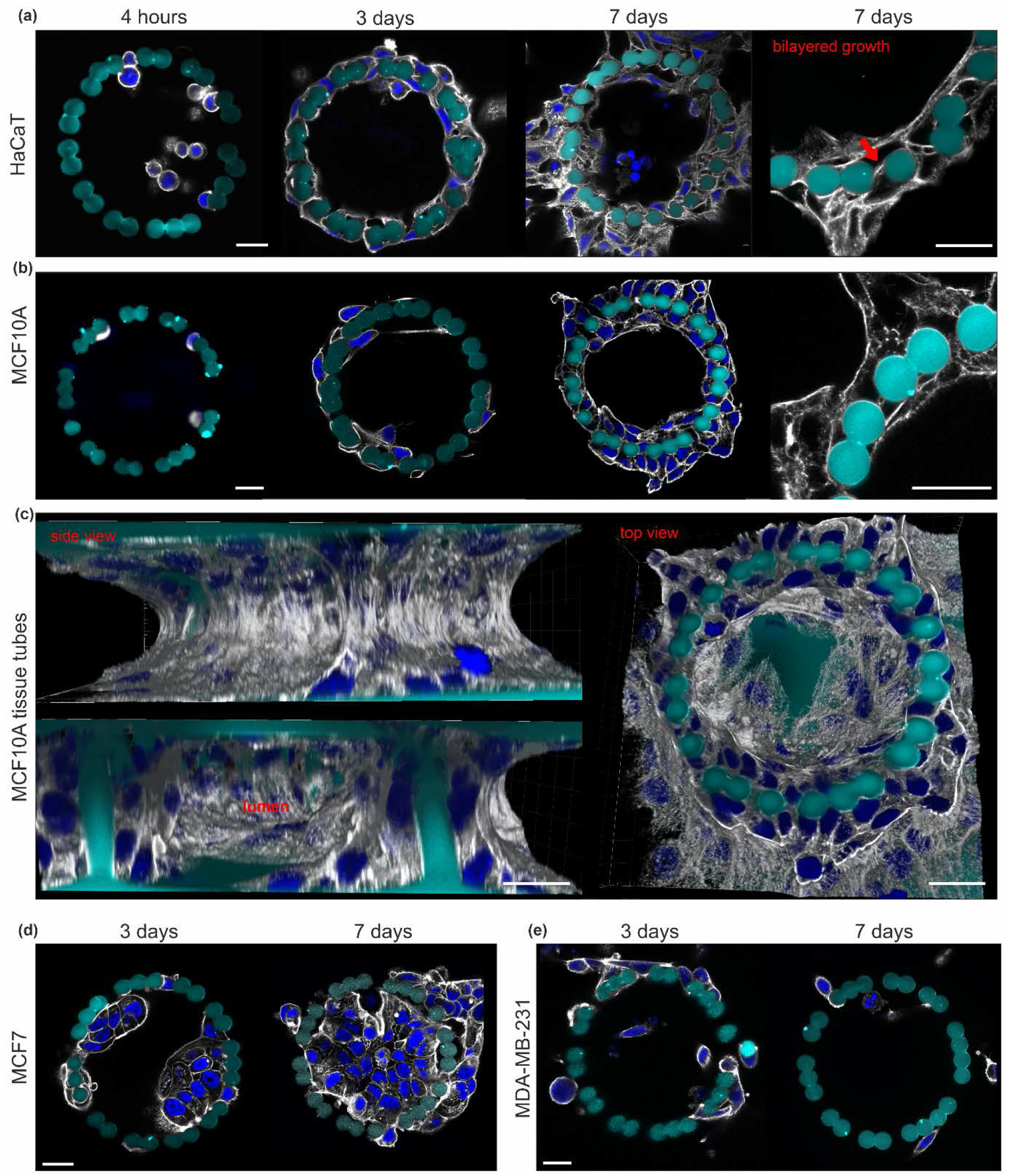
The EPC geometry induced a 3D microtissue growth of normal epithelial cells. Cells were injected into the pillar cages (FN-coated) and cultivated for four hours, three days and seven days. Additional cells were seeded peripherally for supportive media conditioning with soluble signaling molecules. Cells were fixed and stained for the actin cytoskeleton (gray) and nuclei (blue) at 10 μm pillar height. (**a**) and (**b**) HaCaT and MCF10A cell growth over time. Cropped views highlight the bilayered cell cluster morphology (red arrow) around the pillars. (**c**) A 3D reconstructed image stacks show an MCF10A microtissue consisting of 80 pillar-adhered cells (day 7). (**d**) Representative MCF7 tumor cell growth and (**e**) MDA-MB-231 tumor cell dissemination over cultivation time. All cell types were seeded in comparable cell numbers. All scale bars = 20 μm.

In contrast, breast cancer cells showed different growth patterns: Non-invasive MCF7 cells [46] formed bulky aggregates (3 days) that eventually filled the entire pillar cavity (7 days). The lack of cell transmigration through the pillars confirmed a non-invasive growth pattern (Figure 2d). In contrast, MDA-MB-231 cells mirrored their highly-invasive phenotype [47] by continuous pillar transmigration and cell dissemination into the microenvironment (Figure 2D).

These results demonstrated the high biocompatibility of EPCs, suitable for a broad range of long-term 3D cell cultures. The cylindrical pillar topography steered normal epithelial breast and skin cells to form bilayered and lumen-bearing microtissue tubes, whereas monolayered growth was evident on planer substrate areas.

### The EPC geometry modulates actin-mediated cell-cell junctions and matrix adhesion

We analyzed the cytoskeletal organization that could attribute to the distinct monolayered and bilayered growth of breast and skin cells. Actin-mediated cell-cell and cell-matrix contacts were immunostained with the adherence junction (AJ) marker E-cadherin and the focal adhesion (FA) paxillin. Breast and skin cells exhibited a monolayered growth on the planar substrate layer. Both cell types featured condensed E-cadherin signals localized within the actin cortex at cell-cell borders. This result indicated stable cell interconnectivity by adherence junctions (AJ) (Figure 3a, upper rows). Large and partially elongated paxillin spots were frequently localized at the tips of thick actin SFs suggesting the formation of force-transmitting FA-anchored SFs. Such SFs were most abundant in HaCat cells (Figure 3a, lower rows). At the pillar topography, both cell types exhibited a mostly bilayered morphology. In contrast to the monolayered clusters, pillar-bound cells lacked FA-anchored SFs (Figure 3b, lower rows). Paxillin spots were sparsely present within the pronounced cortical actin networks. Figure 3c further illustrates the proximity of paxillin spots to the pillar substrate. Like the monolayered cells, condensed E-cadherin spots were frequently present at cell-cell contact sites indicating AJ-mediated interconnectivity of the bilayered cell tubes (Figure 3b, white arrows).

**Figure 3.**
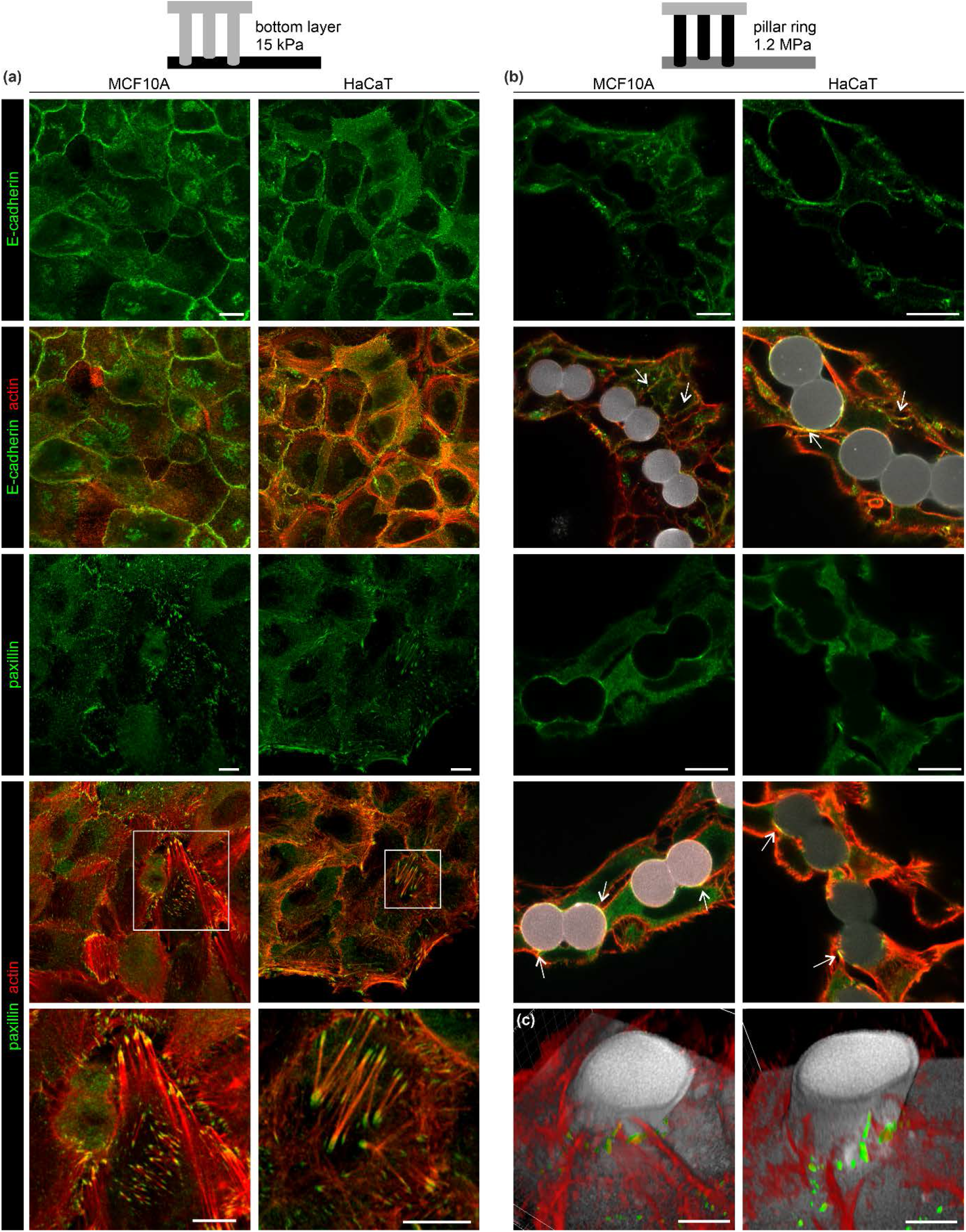
The EPC geometry modulated cell-cell and cell-matrix adhesion. MCF10A and HaCaT were cultivated in EPCs for six days. Cell morphologies were compared at (**a**) the planar bottom layer and (**b**) the pillar ring topography. Cells were fixed and stained for AJs (E-cadherin) and FA complex (paxillin) marker proteins. White arrow heads indicate FA and AJ formation along the pillar surface (gray). Merged images show marker protein and actin cytoskeleton co-localization. Scale bars = 10 μm. (**c**) 3D reconstructions of two independent confocal image stacks. Cross-sections (5 - 10 μm above the bottom substrate) show the spatial proximity of F-actin structures and paxillin spots with the pillar and planar substrate (gray). Scale bars = 5 μm.

This analysis demonstrated that the EPC pillar topography reduced contractile cell-matrix adhesions, typically in monolayered cell clusters. Bilayered breast and skin clusters maintained mechanical stability through cortical actin networks and AJ-mediated interconnectivity.

### Calculation of cell-derived traction force from pillar displacement

We designed the novel EPC array to measure the forces cells exert on the pillar rings. To track individual pillar displacements, fluorescent QDot nanocrystals (10 - 20 nm) were incorporated into the elastomer, forming aggregates of different sizes and shapes (Figure 4a). Pillar displacements were determined utilizing cross-correlation of these patterns [21] To estimate causal force values, we used an analytical approximation based on the Euler-Bernoulli theory of the beam bending model (Figure 4b). For the complete derivation, see Appendix A1. A finite elements method (FEM) simulation for force-mediated pillar bending was performed to evaluate this model. In full agreement with the analytical approximation, FEM revealed asymmetrical displacement fields depending on the height of force application: Pulling at the upper pillar segment shifted the maximum displacement down to the center region (Figure 4c). This shift decreased with force application at the center and lower segments (Figures 4d and e). Force application at the lower part resulted in the largest displacement propagating further to the softer bottom surface plane (Figure 4e). Notably, the displacements retrieved by FEM and numerical approximation showed high coherence: pillar bottom = 79% (0.92 μm / 1.17 μm), pillar center = 97%; (0.7 μm / 0.68 μm) and pillar top = 90% (0.21 μm / 0.19 μm). This evaluation confirmed the bending characteristic of slender beams anchored at both ends to substrates with different stiffness. The high-aspect-ratio pillars showed increasing compliance with distance from their origin. In other words, the effective stiffness decreased with pillar length.

**Figure 4.**
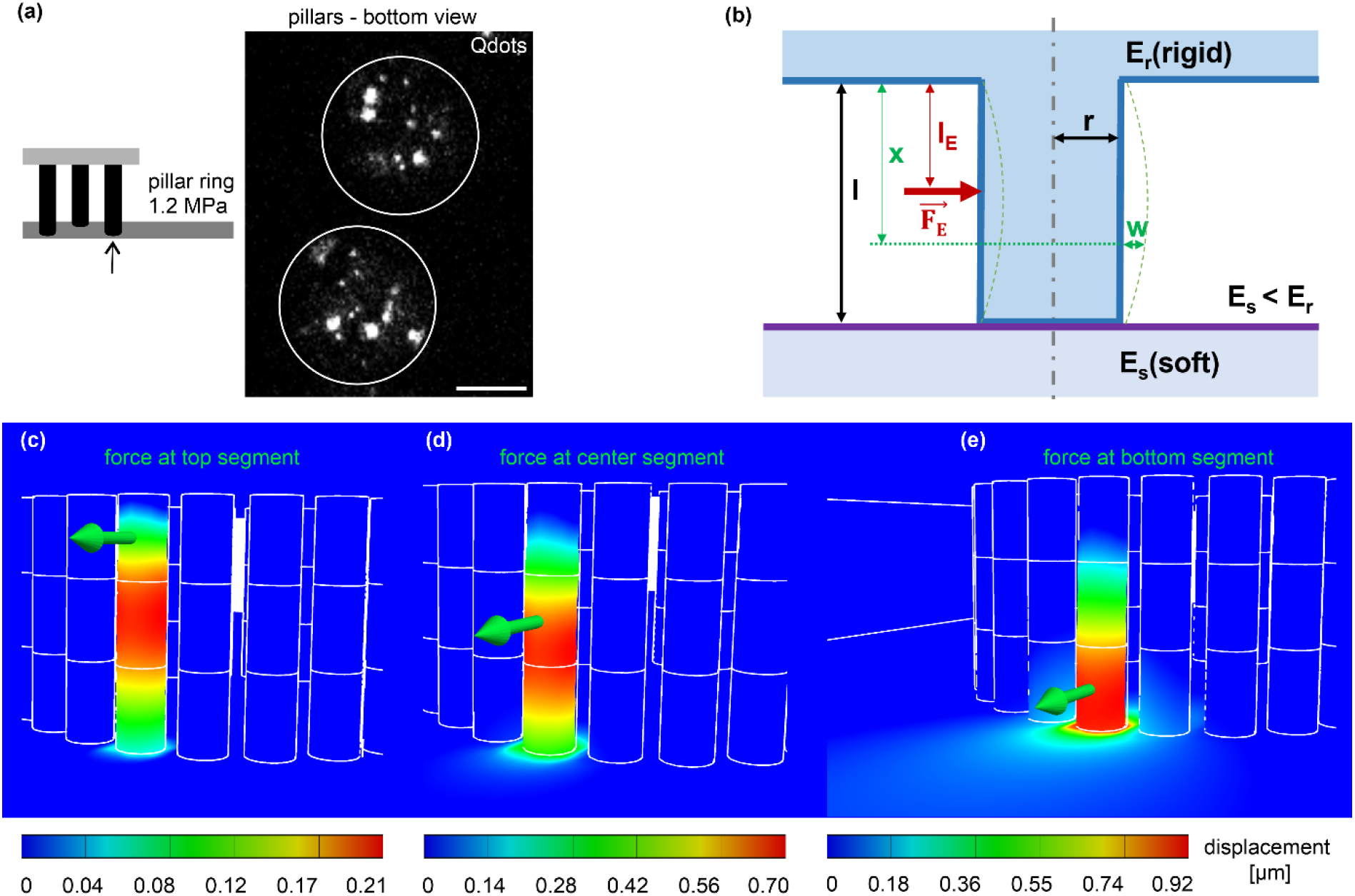
The theoretic framework of cell force calculation from pillar displacement. (**a**) Micrograph shows the cross-section of two adjacent pillars (white outlines) with incorporated fluorescent QDot aggregates to track pillar displacements (confocal image plane: 5 μm distance from the bottom layer). Scale bar = 5 μm. (**b**) Analytic approach for beam bending (blue) with bending parameters used to calculate forces (red). An external single point force F_E_ applied in the distance from the rigid top layer l_E_ (blue) and distance x in which the displacement w was imaged; E_r_ = rigid pillar and the top layer, E_s_ = soft planar bottom layer, l = pillar length and r = pillar radius. (**c - e**) FEM simulations with identical geometrical and material properties for the analytical approximation were used. A representative point force (green arrow) of 189 nN was applied to the top (**c**), center (**d**) and bottom (**e**) segments. Analyzing the FEM simulation results with the analytical approximation (Appendix A1) we received a force of 189 nN.

### Quantification of static and dynamic single-cell forces

To verify assay function, highly contractile myofibroblasts (MFBs) and cardiomyocytes (CMs) were assessed to measure single-cell forces with the EPC approach. Using laser-assisted nanosurgery, a single MFB was cut to measure cell forces from pillar relaxation. A single MFB that spanned between the bottom substrate and pillars was chosen for ablation (Figure 5a, ROI white square). The spatial orientation of five pillars (1 – 5) was analyzed before (t = 0 s) and after cutting (t = 50 s). Figure 5b shows the ROI: MFB (i), was cut (red line), and two adjacent MFBs (ii) remained untreated (Figure 5b). The displacement of QDot marker beads was recorded over the entire pillar volume to create complete pillar bending profiles (Figure 5c). Cell killing resulted in substantial bending of pillars (2, 3, 4 and 5) with a peak at pillar 3, where MFB (i) mainly adhered. Although MFB (i) adhered at the upper pillar region (25 - 46 μm), the highest displacement (0.48 μm) was shifted down to 24 μm (*cf*. Figure 5c, green line, asterisk). A direct comparison of pillar bending peaks and cell adhesion positions highlights the asymmetrical displacement pattern caused by force application at the upper pillar region (cf. Figure 4c and Appendix A1). Of note, occasional QDot tracking errors caused artificial displacements at individual pillar sections, *e*.*g*., cell-free pillar 1. Such technical outliers were excluded from all following force calculations.

**Figure 5.**
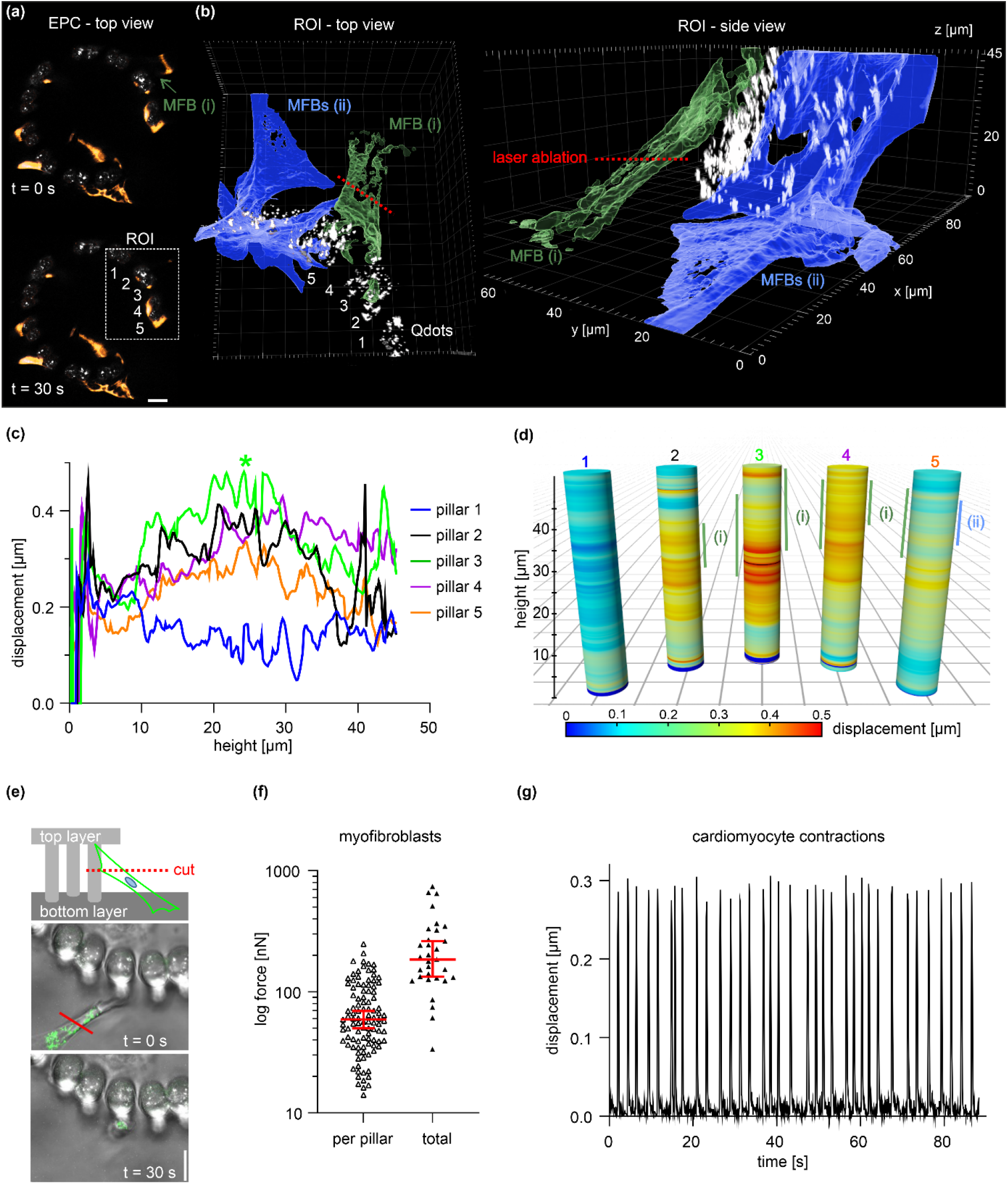
Characterization of pillar bending and quantification of single-cell forces. (**a**) Micrographs display MFB cells (labeled with MitoTracker dye) adhered to a pillar ring (white: QDots). ROI (white square): analyzed pillars before (t = 0 s) and after cell cut (t = 50 s) with a nanosecond-pulsed ablation laser. Scale bar = 20 μm. For a complete image series of cell ablation, see Movie S2. (**b**) A surface-rendered 3D reconstruction of a confocal image stack highlights the spatial orientation of MFB (i) (green) and two adjacent untreated MFBs (ii) (blue) before ablation (t = 0 s). (**c**) Displacements were measured along entire pillar heights. Confocal image stacks of the five analyzed pillars in the ROI (a) were taken at t = 0 s and t = 50 s. Green asterisk: max. displacement peak. (**d**) Graphical representation of (c) compares pillar bending profiles with cell adhesion sites (vertical bars). (**e**) Experimental setup for quantitative force analysis of single MFBs that connected the bottom substrate and pillars. Images show a representative MFB (MitoTracker dye and brightfield) pre (t = 0 s) and post-cut (t = 30 s). Scale bar = 10 μm. (**f**) The plot summarizes the forces cells exerted on individual pillars (n = 103, three independent samples) and the corresponding sum of forces exerted per cell (n = 31). Horizontal line: median; whiskers: 95% CI. (**g**) The plot shows the pillar displacement retrieved from a spontaneously contracting cardiomyocyte. For the complete timelapse series, see Movie S3.

Next, we quantified the force generation of MFBs using the same experimental setup (Figure 5e). Pillar displacements were measured at the imaging plane of cell ablation. To circumvent determining the actual substrate adhesion position for each cell, a fixed force application point at 5 μm pillar height was assumed (that corresponded to l_E_ = 45 μm) (see Appendix A1). Depending on the actual height of force application, this approach slightly underestimated absolute force values (cf. Figure 4e). After cell ablation, pillar relaxations typically reached a plateau phase at t ≥ 30 s (Figure S1). This time point was applied for all following force calculations. For single MFBs, we calculated a median force per pillar of 73 nN. Since cells often adhered to several pillars (*cf*. Figure 5b), for those cells, the median total cell force was substantially higher (227 nN) (Figure 5f). In addition, we tested to resolve dynamic forces by using beating cardiomyocytes (CMs) (Figure 5f). We measured a periodic contraction of 0.4 Hz and a mean cell force per pillar of 26 nN (n = 60 pillars). These validation experiments demonstrated that the EPC force approach resolved static and fast dynamic cell forces.

### Quantitation of epithelial microtissue forces

EPC-derived microtissues showed monolayered and bilayered cell areas accompanied by different cytoskeleton organizations (cf. Figures 2 and 3). To link these morphological alterations with modulated cell contractility, comparative force measurements were performed with cells on a planar substrate and pillar topography. Bilayered cell tubes were entirely detached from pillars by rigor circular cuts (Figures 6a and b). Displacements were measured at 5 μm pillar height, where maximum deflections were assumed (cf. Figure 4e). However, only neglectable pillar relaxation was observed, similar to the cell-free pillar control. This result contrasted the substantial pillar displacement of the MFB control (Figure 6c). A cumulative force analysis confirmed the small contractility of breast and skin tissue tubes (Figure 6d). In detail, the mean sum of contractile forces of cut MCF10A (10 nN) and HaCaT (15 nN) cell tubes was insignificantly different from the untreated controls (MCF10A ctrl: 8 nN; HaCaT ctrl: 10 nN). In contrast, single CMs and MFBs exerted mean force amplitudes between 26 nN and 73 nN that were up to seven-fold higher compared to multicellular MCF10A tubes (strong effect size Cohen’s d = 2.9, p < 0.0001). Nevertheless, HaCat-derived tubes showed a significantly higher force (+32%) compared to MCF10A breast cell tubes (moderate effect size Cohen’s d = 0.6, p < 0.0001). Notably, MCF10A force was insignificant from the cell-free control (11 nN). In addition, MCF10A microtissue treatment with lysophosphatidic acid (LPA) led to a temporal increase of total force (+73%, 17.5 nN at 40 min.) and hence confirmed the inducibility of actomyosin contractibility in MCF10A cell tubes (Figure 6d). These results suggested that pillar-bound microtissues of breast and skin cells maintained a relatively inactive actomyosin apparatus.

**Figure 6.**
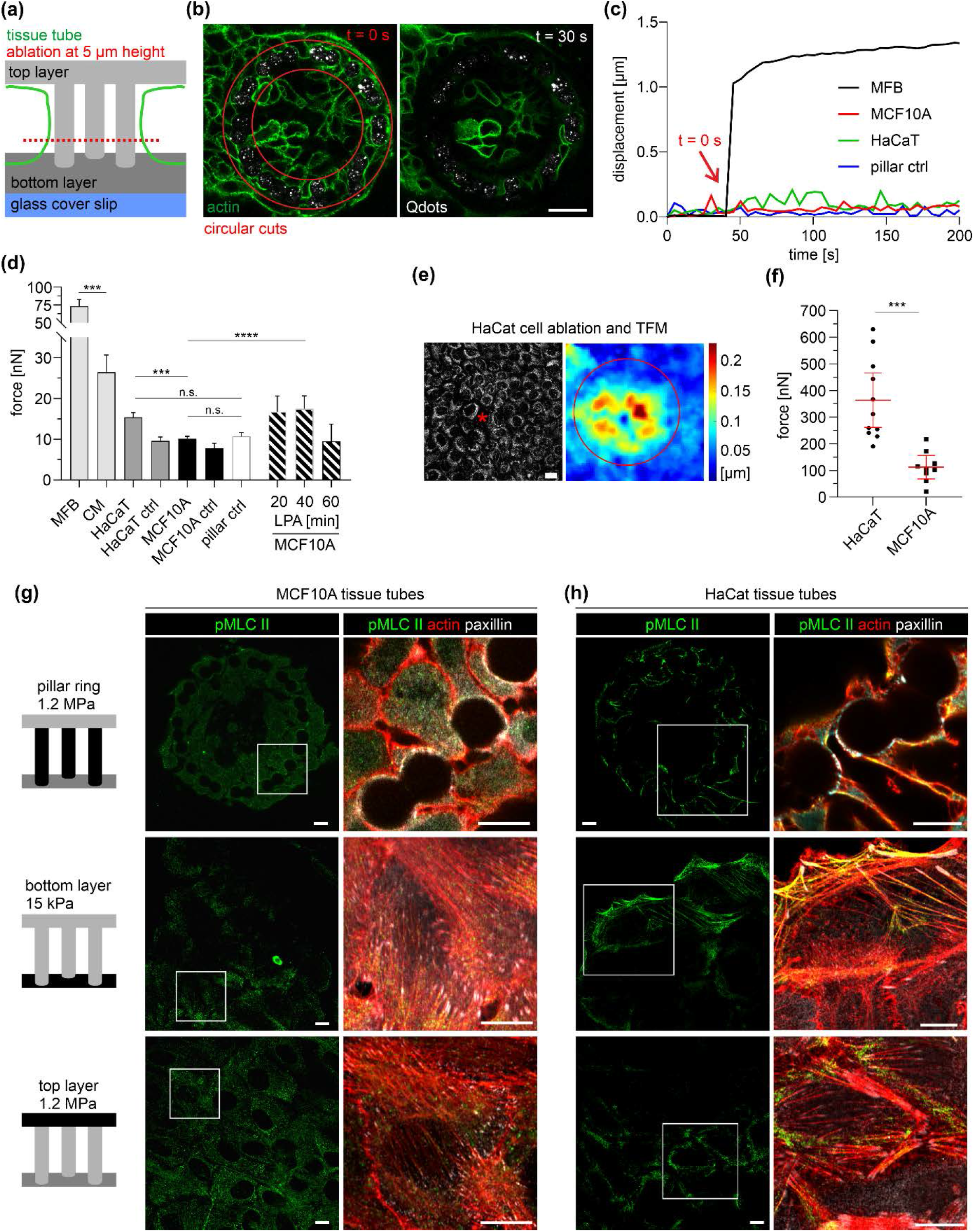
Force quantification in mono- and bilayered microtissues. (**a**) Experiment design of cell ablation for cell force retrieval. (**b**) Micrograph shows a representative MCF10A cell tube (green, lifeActRFP) before (t = 0 s) and after (t = 30 s) pillar detachment (white spots, QDot marker beads) by circular cuts (red outlines). Scale bar = 20 μm. (**c**) The plot compares the displacements of single pillars after cell cutting (red arrow) of a single MFB, HaCaT and MCF10A cell tubes and a cell-free pillar (pillar ctrl). (**d**) Overall comparison of the sum of contractile forces calculated from total pillar displacements. Bars: mean, whiskers: 95% CI. At least three independent experiments were performed: MCF10A (ablated): n = 20 tubes, total pillars n = 407; MCF10A ctrl (untreated): n = 5 tubes, total pillars n = 92; HaCaT (ablated): n = 15 tubes, total pillars n= 338; HaCaT (untreated): n = 7 tubes, total pillars n= 131; pillar ctrl: n= 84 and LPA treatment (30 μM) (n = 1 tube), total pillars: n = 14 (20 min), n = 20 (40 min), n = 1 (60 min). (**e**) 2D TFM with cells on planar substrates (15 kPa). A representative deformation field is shown, yielded from single cell ablation (asterisk) of a HaCaT monolayer labeled by MitoTracker dye. Red outline: ROI for force calculation. (**f**) A scatter plot shows the sum of contractile force after single-cell ablation in (HaCaT (n = 10) and MCF10A (n = 9) monolayers. Horizontal line: mean; whiskers: 95% CI. (**g**) and (**h**) Micrographs show the actomyosin cytoskeleton of EPC-derived microtissues. Cells were fixed and immunostained against activated myosin II (pMLC II, green), FA-protein paxillin (gray) and actin (red). Pillar-bound cells were imaged at 10 μm height. For a single-channel display, see Figure S2. Scale bars = 10 μm; n.s.: p ≥ 0.05; ***: p < 0.001; ****: p < 0.0001).

In contrast, single-cell ablation of monolayered clusters on planar 15 kPa substrates resulted in substantial deformation fields that circularly propagated from the killed cell (Figure 6e). The mean sum of contractile force was substantially increased by 25-fold for HaCaT (364 nN) and 11-fold for MCF10A (112 nN) (Figure 6f), compared with the pillar-bound clusters. Finally, we compared the measured cell contractility with the cytoskeleton organization of mono- and bilayered cell clusters by staining for phosphorylated myosin II (pMLC II) (Figure 6). At the planar top and bottom layers, MCF10A and HaCat monolayers pMLC II localized at large SF bundles anchored to FAs. These force-transmitting actomyosin bundles appeared more pronounced in HaCat than in MCF10A. In contrast, bilayered clusters lacked such actomyosin SF formation. However, occasional pMLC II spots were located within the actin cortex of HaCaT cells (Figure 6h) and absent in MCF10A (Figure 6g). These stainings fit the generally high monolayer contractility and the highest amplitudes for HaCaT cells (cf. Figure 6d).

Together, the force analyses indicated that bilayered cell morphologies were accompanied by reduced actomyosin contractability resulting in low-tensional breast and skin microtissues. Notably, such low traction forces were reproducibly detected by our method.

## Discussion

Mechanical properties of tissues, such as stiffness [9], topography [10] and geometry [11], regulate cell contractility. Cell forces, in turn, modulate cell shape [2] and differentiation [3]. Investigating stiffness and topography-regulated force generation in 3D epithelial microtissues remained technically challenging. To this end, we developed a new quantitative cell force approach to simultaneously analyze the impact of substrate stiffness and topographies on tissue geometry and cell contractility of epithelial microtissues.

We showed that microfabricated EPC arrays were suitable for cultivating tumorous and normal epithelial, stromal, and cardiac cell types. The array incorporated large diffusion channels facilitating continuous nutrition exchange for excellent cell viability and homogenous surface functionalization with ECM proteins. The array consisted of 40 spatially separated EPC samples to perform multiposition experiments with statistical relevance. The optical clarity of PDMS silicone rubber [48] enabled confocal fluorescent microscopy analyses of even 50 μm thick 3D microtissues in high spatial and temporal resolution. We used the unique EPC geometry to engineer bilayered cell tubes derived from human breast and skin cell lines. These contact-inhibited microtissues formed within seven days and maintained a homeostatic tissue state.

In contrast, MDA-MB231 breast cancer cells resembled their highly invasive phenotype with transmigration through the pillar gaps and dissemination into the microenvironment [49]. MCF7 cells proliferated to dense cell aggregates within the pillar cage boundary. A comparable growth pattern has been described in a study that used circular pillar arrangements to estimate expansive forces of growing tumor cell spheroids [50]. These findings confirmed the feasibility of EPCs in analyzing cell growth and migration of a wide range of epithelial cells. Moreover, we suggest that the EPC arrays are adaptable to a broad range of microtissue engineering approaches. The present work used EPCs that consisted of stiff (1.2 MPa) high aspect ratio pillars with a seamless transition to the planar bottom layer (15 kPa). The modular EPC design allows modulating the stiffness gradient, *e*.*g*., by using softer (< 15 kPa) bottom layer substrates. Of note, there is a technical limit for ultrasoft elasticities (< 1 kPa) since pillar molds would partially sink into such compliant substrates. Nevertheless, the implemented stiffness gradient enables to study durotactic phenomenons within a defined 3D topography and thus complements stiffness-driven cell migration and polarization studies on 2D elastomeric substrates [51, 52].

However, the present work focused on epithelial microtissue engineering. The EPC array was designed to measure force microscopy in a mixed geometry of planar surfaces and cylindrical pillar topography. This approach differs from well-established approaches, using dense arrays of free-standing micropillars on which cells adhere. Cell forces were usually calculated from the deflection of pillars exhibiting a simple and well-understood geometry [26, 28, 53, 54]. For our approach, we needed to develop an analytical approximation to describe the bending of slender beams, whose ends are connected to planar substrates of different stiffness (see Appendix A1). FEM simulations confirmed the results of this approximation that based on the Euler-Bernoulli theory of beam bending. Moreover, the theoretical prediction was further supported by the experiments on highly contractile myofibroblasts where we observed asymmetric bending profiles. These cells exerted substantial traction force (227 nN) on the EPCs. This force was in the range of heart muscle cell contractility measured by the deflection of free-standing micropillars (140 nN – 400 nN) [42]. Moreover, the analysis of beating cardiomyocytes resolved fast cell contractions (0.4 Hz) that were comparable to previous 2D TFM measurements (0.6 Hz) [35] and with a force amplitude (26.4 nN) as described for an approach of 3D microprinted elastic wheel-like scaffolds (50 nN) [54]. These findings verified that our method could resolve both static and dynamic forces of single cells.

We aimed to investigate the force generation of 3D skin and breast epithelium microtissues. Indeed, we found fundamental changes in tissue shape, cytoskeletal organization and actomyosin contractility, which depended on substrate stiffness and topography. On planar substrate areas, skin and breast cells formed monolayered clusters with spread cell shapes and abundant actin SF and FA formation that further increased with substrate stiffness. Consequently, the highest single-cell force (343 nN) was accompanied by abundant actomyosin SFs and the large FA patches in HaCaT monolayers. MCF10A single-cell forces (182 nN) were in line with a comparable force study on 10 kPa planar elastomeric substrates (150 nN) [55]. Together, our findings confirmed the well-described mechanoresponse of 2D epithelial cell cultures to substrate stiffness, leading to increased actomyosin contractility mediated by FA-bound SF bundles [13, 56-58].

More surprisingly, the pillar rings induced the growth of AJ-interconnected bilayered microtissues, consisting of roundish cells with pronounced cortical actin networks. These SF-lacking cell morphology resembled organotypic cultures for HaCaT skin equivalents and MCF10A breast gland spheroids. Indeed, EPC-derived microtissues were rather simplified versions of skin equivalents and breast gland acini due to the lack of stratification [59, 60] or basoapical polarization [61, 62]. Nevertheless, the topography of circularly arranged pillars induced a mechanoadaptation of HaCaT and MCF10A cells towards more physiologic tissue morphologies.

Compared to their monolayered counterparts, bilayered clusters generated only marginal total tissue forces (10 - 15 nN). For multicellular MCF10A, tube contractility was undistinguishable from the cell-free pillar control (≤ 11 nN). Suppose one assumes that each cell within a cell tube contributed equally to the total tissue tension. In that case, single-cell forces can at least be roughly approximated: Based on the count of 80 pillar-bound cells, a typical MCF10A microtissue consisted of (*cf*. Figure 2c), a mean single cell force of results in a value of 0.22 nN. Notably, such piconewton-scaled single-cell forces were far below the technical detection limit of pillar displacements.

Nevertheless, even the measured total force of EPC-derived MCF10A cell tubes contrasted the comparably high actomyosin contractility (20 nN) described for single MCF10A cells exhibiting an actin SF-rich cytoskeleton [63]. Our findings linked the utmost relaxed states of EPC-derived breast and skin microtissues to the lack of actomyosin SF contractility. However, we found cell contractility in MCF10A cell tubes was inducible by activating Rho/Rock-signaling upon LPA treatment [64]. Together these results suggested that the functional contractile apparatus of pillar-bound cells actively maintained the observed low tensional tissue state.

At first glance, these low-tensional states appeared inconsistent with the high material stiffness of pillars (1.2 MPa) at which cells adhered: the same cells exerted substantial actomyosin contractility on softer substrates (15 kPa). Moreover, previous works showed that comparable substrate stiffness (12 kPa) triggered EMT-like cytoskeletal reorganization and actomyosin contractility in SF-lacking MCF10A spheroids [34, 65]. These findings raised the question of how the EPC topography modulated cellular mechanosensation: Micropillars are generally defined by substrate surface elasticity (local stiffness) and structural compliance (effective stiffness). The high aspect ratio of the EPC topography generated a low effective pillar stiffness sensed by pillar-bound cells. This topographical cue was cellularly transduced into reduced actomyosin contractility. A comparable topography effect on cell contractility has been described for hydrogel-embedded interstitial cells anchored between two elastic posts [66, 67]. In addition, micron-scaled confinements with pillar diameters (5 – 10 μm) and spacing (5 – 10 μm), comparable to the EPCs, affected the cell shape and migration of single fibroblasts by modifying cell-matrix adhesion and actomyosin contractility [10]. These findings suggested a topography-driven mechanoadaptation that overwrote local substrate stiffness sensing, causing low tensional states of breast and skin microtissues. Interestingly, a certain contractility (15 nN) was evident in HaCaT cell tubes and associated with actomyosin activity within the actin cortex. This finding implicated that cell tube tension could have originated from the cortical actin cytoskeleton [68]. Elegant work supports this explanation by demonstrating the embedment of SFs within the actin cortex of epithelial cells and its contribution to actomyosin contractility [69]. Besides the present work, comparable quantification of cortical traction forces in bilayered epithelia has not been reported. Our findings show a cellular mechanoadaptation to the EPC topography that modulated actomyosin contractility. Thereby bilayered breast and skin cells actively maintained a homeostatic tissue state with low cortical tension.

## Conclusions

Cell contractility is an essential modulator of tissue shape and function. Cellular forces broadly spread depending on cell type and the culturing conditions [70]. Meaningful microtissue formation and homeostasis studies need, thus, a better understanding of cell force modulation by mechanical microenvironmental cues. We introduced a novel 3D cell culture device to explore the mechanobiological regulation of cell shape and function in response to substrate stiffness and topography. The EPC array enabled quantifying and comparing cell contractility within defined mono- and bilayered epithelial microtissues. Our approach bridges stiffness and topographical cues with cell force microscopy of engineered 3D microtissues. The EPC arrays could be helpful to a wide range of scientific questions that address mechanobiological regulation circuits in microtissue development and homeostasis.

## Supplementary Materials

**Figure S1:** Time-dependent pillar relaxation., **Figure S2:** Modulation of the actomyosin cytoskeleton by the EPC geometry, **Movie S1:** Lateral cell injection of MCF10A cells using a glass microcapillary, **Movie S2:** Laser ablation (white arrowhead) of a myofibroblast (labeled by MitoTracker dye). Pillars with incorporated QDots (gray) were used for displacement tracking, **Movie S3:** Time lapse of spontaneous beating cardiomyocytes (labeled by MitoTracker dye, orange). Pillars with incorporated QDots (green) were used for displacement tracking (Imaging interval: 90 ms); **Appendix A1**: Analytical approximation of EPC-derived cell forces.

## Author Contributions

Conceptualization, E.N., R.M., B.H.; Supervision, E.N., B.H., R.M.; funding acquisition, R.M.; methodology, R.M., G.D., R.S., N.H., J.K.; formal analysis, L.E., G.D., R.S., E.N.; visualization, L.E, R.S., L.L., J.K., E.N.; writing—original draft, R.M. L.E., E.N.; writing—review & editing, E.N., B.H., R.M. All authors have read and agreed to the published version of the manuscript.

## Funding

Funded by the Deutsche Forschungsgesellschaft (DFG, German Research Foundation) 363055819/GRK2415 through SPP1782 within the projects H02384/2 and ME1458/8.

## Institutional Review Board Statement

Not applicable.

## Informed Consent Statement

Not applicable.

## Data Availability Statement

The datasets supporting the conclusions of this article are either available within the paper and its supplementary information files or from the corresponding author upon reasonable request.

## Acknowledgments

The authors would like to thank Nils Hersch for his skilled laboratory work.

## Conflicts of Interest

The authors declare no conflict of interest.

## Abbreviations

CI: (confidence interval)
ECM: (extracellular matrix)
FA: (focal adhesion)
AJ: (adherence junction)
pMLC II: (phosphorylated myosin light chain II)
SF: (stress fiber)
TFM: traction force microscopy
EPC: (elastomeric pillar cage)
MFB: (myofibroblast)
CM: (cardiomyocyte)

## Notes

### Competing Interest Statement

The authors have declared no competing interest.

## References

1. Butcher, D.T., T. Alliston, and V.M. Weaver, A tense situation: forcing tumour progression. Nat Rev Cancer, 2009. 9(2): p. 108–22.

2. McBeath, R., et al., Cell shape, cytoskeletal tension, and RhoA regulate stem cell lineage commitment. Dev Cell, 2004. 6(4): p. 483–95.

3. Kilian, K.A., et al., Geometric cues for directing the differentiation of mesenchymal stem cells. Proc Natl Acad Sci U S A, 2010. 107(11): p. 4872–7.

4. Rosowski, K.A., et al., Edges of human embryonic stem cell colonies display distinct mechanical properties and differentiation potential. Sci Rep, 2015. 5: p. 14218.

5. Gjorevski, N. and C.M. Nelson, Mapping of mechanical strains and stresses around quiescent engineered three-dimensional epithelial tissues. Biophys J, 2012. 103(1): p. 152–62.

6. Nelson, C.M., et al., Emergent patterns of growth controlled by multicellular form and mechanics. Proc Natl Acad Sci U S A, 2005. 102(33): p. 11594–9.

7. Gomez, E.W., et al., Tissue geometry patterns epithelial-mesenchymal transition via intercellular mechanotransduction. J Cell Biochem, 2010. 110(1): p. 44–51.

8. Inman, J.L., et al., Mammary gland development: cell fate specification, stem cells and the microenvironment. Development, 2015. 142(6): p. 1028–42.

9. Saez, A., et al., Rigidity-driven growth and migration of epithelial cells on microstructured anisotropic substrates. Proc Natl Acad Sci U S A, 2007. 104(20): p. 8281–6.

10. Ghibaudo, M., et al., Substrate topography induces a crossover from 2D to 3D behavior in fibroblast migration. Biophys J, 2009. 97(1): p. 357–68.

11. Thery, M., et al., Cell distribution of stress fibres in response to the geometry of the adhesive environment. Cell Motil Cytoskeleton, 2006. 63(6): p. 341–55.

12. Geiger, B., et al., Transmembrane crosstalk between the extracellular matrix--cytoskeleton crosstalk. Nat Rev Mol Cell Biol, 2001. 2(11): p. 793–805.

13. Geiger, B., J.P. Spatz, and A.D. Bershadsky, Environmental sensing through focal adhesions. Nat Rev Mol Cell Biol, 2009. 10(1): p. 21–33.

14. Cai, Y., et al., Cytoskeletal coherence requires myosin-IIA contractility. J Cell Sci, 2010. 123(Pt 3): p. 413–23.

15. Hu, S., et al., Intracellular stress tomography reveals stress focusing and structural anisotropy in cytoskeleton of living cells. Am J Physiol Cell Physiol, 2003. 285(5): p. C1082–90.

16. Vasquez, C.G. and A.C. Martin, Force transmission in epithelial tissues. Dev Dyn, 2016. 245(3): p. 361–71.

17. Chen, C.S., et al., Geometric control of cell life and death. Science, 1997. 276(5317): p. 1425–8.

18. Yu, S.M., et al., Mechanical Adaptations of Epithelial Cells on Various Protruded Convex Geometries. Cells, 2020. 9(i6).

19. Bao, M., et al., 3D microniches reveal the importance of cell size and shape. Nat Commun, 2017. 8(1): p. 1962.

20. Dembo, M. and Y.L. Wang, Stresses at the cell-to-substrate interface during locomotion of fibroblasts. Biophys J, 1999. 76(4): p. 2307–16.

21. Merkel, R., et al., Cell force microscopy on elastic layers of finite thickness. Biophys J, 2007. 93(9): p. 3314–23.

22. Butler, J.P., et al., Traction fields, moments, and strain energy that cells exert on their surroundings. Am J Physiol Cell Physiol, 2002. 282(3): p. C595–605.

23. Angelini, T.E., et al., Cell migration driven by cooperative substrate deformation patterns. Phys Rev Lett, 2010. 104(16): p. 168104.

24. Trepat, X., et al., Physical forces during collective cell migration. Nature Physics, 2009. 5(6): p. 426–430.

25. Tambe, D.T., et al., Collective cell guidance by cooperative intercellular forces. Nat Mater, 2011. 10(6): p. 469–75.

26. Tan, J.L., et al., Cells lying on a bed of microneedles: an approach to isolate mechanical force. Proc Natl Acad Sci U S A, 2003. 100(4): p. 1484–9.

27. du Roure, O., et al., Force mapping in epithelial cell migration. Proceedings of the National Academy of Sciences of the United States of America, 2005. 102(7): p. 2390.

28. Ghassemi, S., et al., Cells test substrate rigidity by local contractions on submicrometer pillars. Proc Natl Acad Sci U S A, 2012. 109(14): p. 5328–33.

29. Serrano, R., et al., Three-Dimensional Monolayer Stress Microscopy. Biophys J, 2019. 117(1): p. 111–128.

30. Legant, W.R., et al., Measurement of mechanical tractions exerted by cells in three-dimensional matrices. Nat Methods, 2010. 7(12): p. 969–71.

31. Hall, M.S., et al., Fibrous nonlinear elasticity enables positive mechanical feedback between cells and ECMs. Proc Natl Acad Sci U S A, 2016. 113(49): p. 14043–14048.

32. Mark, C., et al., Collective forces of tumor spheroids in three-dimensional biopolymer networks. Elife, 2020. 9.

33. Ulbricht, A., et al., Cellular mechanotransduction relies on tension-induced and chaperone-assisted autophagy. Curr Biol, 2013. 23(5): p. 430–5.

34. Eschenbruch, J., et al., From Microspikes to Stress Fibers: Actin Remodeling in Breast Acini Drives Myosin II-Mediated Basement Membrane Invasion. Cells, 2021. 10(8).

35. Hersch, N., et al., The constant beat: cardiomyocytes adapt their forces by equal contraction upon environmental stiffening. Biol Open, 2013. 2(3): p. 351–61.

36. Ahrens, D., et al., A Combined AFM and Lateral Stretch Device Enables Microindentation Analyses of Living Cells at High Strains. Methods Protoc, 2019. 2(2).

37. Houben, S., N. Kirchgeßner, and R. Merkel, Estimating force fields of living cells - Comparison of several regularization schemes combined with automatic parameter choice, in Lecture Notes in Computer Science (including subseries Lecture Notes in Artificial Intelligence and Lecture Notes in Bioinformatics). 2010.

38. Boussinesq, J., Application des potentiels à l’étude de l’équilibre et du mouvement des solides élastiques. 1885, Paris, F: Gauthier-Villars.

39. Landau, L.D. and E.M. Lifshitz, Course of Theoretical Physics. VII Theory of Elasticity. 1986, Oxford, UK: Pergamon Press.

40. Maloney, J.M., et al., Influence of finite thickness and stiffness on cellular adhesion-induced deformation of compliant substrata. Physical Review E, 2008. 78: p. 041923.

41. Cesa, C.M., et al., Micropatterned silicon elastomer for high resolution analysis of cell force patterns. Reviews of Scientific Instruments, 2007. 78: p. 034301.

42. Kajzar, A., et al., Toward Physiological Conditions for Cell Analyses: Forces of Heart Muscle Cells Suspended Between Elastic Micropillars. Biophysical Journal, 2008. 94(5): p. 1854–1866.

43. Timoshenko, S.P. and J.N. Goodier, Theory of Elasticity. 1970, Auckland, NZ: McGraw-Hill.

44. Boukamp, P., et al., Normal keratinization in a spontaneously immortalized aneuploid human keratinocyte cell line. The Journal of Cell Biology, 1988. 106(3): p. 761.

45. Soule, H.D., et al., Isolation and Characterization of a Spontaneously Immortalized Human Breast Epithelial Cell Line, MCF-10. Cancer Research, 1990. 50(18): p. 6075.

46. Soule, H.D., et al., A Human Cell Line From a Pleural Effusion Derived From a Breast Carcinoma2. JNCI: Journal of the National Cancer Institute, 1973. 51(5): p. 1409–1416.

47. Brinkley, B.R., et al., Variations in Cell Form and Cytoskeleton in Human Breast Carcinoma Cells *in Vitro*. Cancer Research, 1980. 40(9): p. 3118.

48. Piruska, A., et al., The autofluorescence of plastic materials and chips measured under laser irradiation. Lab Chip, 2005. 5(12): p. 1348–54.

49. Kenny, P.A., et al., The morphologies of breast cancer cell lines in three-dimensional assays correlate with their profiles of gene expression. Molecular Oncology, 2007. 1(1): p. 84–96.

50. Aoun, L., et al., Measure and characterization of the forces exerted by growing multicellular spheroids using microdevice arrays. PLoS One, 2019. 14(5): p. e0217227.

51. Wormer, D.B., et al., The focal adhesion-localized CdGAP regulates matrix rigidity sensing and durotaxis. PLoS One, 2014. 9(3): p. e91815.

52. DuChez, B.J., et al., Durotaxis by Human Cancer Cells. Biophys J, 2019. 116(4): p. 670–683.

53. du Roure, O., et al., Force mapping in epithelial cell migration. Proc Natl Acad Sci U S A, 2005. 102(7): p. 2390–5.

54. Klein, F., et al., Elastic Fully Three-dimensional Microstructure Scaffolds for Cell Force Measurements. Advanced Materials, 2010. 22(8): p. 868–871.

55. Kraning-Rush, C.M., J.P. Califano, and C.A. Reinhart-King, Cellular traction stresses increase with increasing metastatic potential. PloS one, 2012. 7(2): p. e32572–e32572.

56. Chrzanowska-Wodnicka, M. and K. Burridge, Rho-stimulated contractility drives the formation of stress fibers and focal adhesions. J Cell Biol, 1996. 133(6): p. 1403–15.

57. Discher, D.E., P. Janmey, and Y.L. Wang, Tissue cells feel and respond to the stiffness of their substrate. Science, 2005. 310(5751): p. 1139–43.

58. Gupta, M., et al., Adaptive rheology and ordering of cell cytoskeleton govern matrix rigidity sensing. Nat Commun, 2015. 6: p. 7525.

59. Weinmuellner, R., et al., Long-term exposure of immortalized keratinocytes to arsenic induces EMT, impairs differentiation in organotypic skin models and mimics aspects of human skin derangements. Archives of Toxicology, 2018. 92(1): p. 181–194.

60. Maas-Szabowski, N., A. Starker, and N.E. Fusenig, Epidermal tissue regeneration and stromal interaction in HaCaT cells is initiated by TGF-alpha. J Cell Sci, 2003. 116(Pt 14): p. 2937–48.

61. Debnath, J., S.K. Muthuswamy, and J.S. Brugge, Morphogenesis and oncogenesis of MCF-10A mammary epithelial acini grown in three-dimensional basement membrane cultures. Methods, 2003. 30(3): p. 256–268.

62. Gaiko-Shcherbak, A., et al., The Acinar Cage: Basement Membranes Determine Molecule Exchange and Mechanical Stability of Human Breast Cell Acini. PLOS ONE, 2015. 10(12): p. e0145174.

63. Lemmon, C.A., et al., Shear force at the cell-matrix interface: enhanced analysis for microfabricated post array detectors. Mechanics & chemistry of biosystems : MCB, 2005. 2(1): p. 1–16.

64. Ridley, A.J. and A. Hall, The small GTP-binding protein rho regulates the assembly of focal adhesions and actin stress fibers in response to growth factors. Cell, 1992. 70(3): p. 389–99.

65. Gaiko-Shcherbak, A., et al., Cell Force-Driven Basement Membrane Disruption Fuels EGF- and Stiffness-Induced Invasive Cell Dissemination from Benign Breast Gland Acini. Int J Mol Sci, 2021. 22(8).

66. John, J., et al., Boundary stiffness regulates fibroblast behavior in collagen gels. Ann Biomed Eng, 2010. 38(3): p. 658–73.

67. Legant, W.R., et al., Microfabricated tissue gauges to measure and manipulate forces from 3D microtissues. Proc Natl Acad Sci U S A, 2009. 106(25): p. 10097–102.

68. Chugh, P., et al., Actin cortex architecture regulates cell surface tension. Nat Cell Biol, 2017. 19(6): p. 689–697.

69. Vignaud, T., et al., Stress fibres are embedded in a contractile cortical network. Nat Mater, 2021. 20(3): p. 410–420.

70. Eyckmans, J. and C.S. Chen, 3D culture models of tissues under tension. J Cell Sci, 2017. 130(1): p. 63–70.

