## Appendix A1 for "Elastomeric Pillar Cages Modulate Actomyosin Contractility of Epithelial Microtissues by Substrate Stiffness and Topography"

### Supplementary Material Appendix A1: Compliance of a Beam Connecting Two Elastic Half Spaces

R. Merkel

#### 1 Introduction

As described in the main text of this article, we studied cells within a circular array of elastic micropillars connected to a compliant substrate. With confocal microscopy we were able to observe displacements of fluorescent marker particles embedded in the micropillar matrix. These displacements indicated active cell forces that we searched to quantify. Unfortunately the mechanical layout of this system differs from the classical textbook examples of beam bending. Because both ends of each micropillar transmit forces and moments to the respective flat substrates this beam is statically indetermined. Moreover, its ends are connected in a flexible way. Therefore standard boundary conditions (fixed ends or fixed angles at the ends) do not apply and special treatment is necessary. As always, a fully numerical solution of this mechanical problem is easily accessible by the well-established finite element method (FEM), however, this would require time intensive calculations for each specific load case. Moreover, due to the many control parameters entering the calculation (e.g. beam diameter and length, material stiffnesses, or attachment geometries of cells) it is very difficult to get an overview over the structure of the general mechanical problem. Therefore we decided to use an analytical approach based on the Euler-Bernoulli theory of beam bending (see any advanced textbook of technical or continuum mechanics, e.g. [39,43]).

#### 2 Basic Equations

Our approach is based on the following approximation. We neglect cross talk between different pillars and assume perfect cylindrical shape for each cylinder. Therefore we study the behavior of a cylindrical micropillar of length $l$ and radius $r$. It protrudes vertically from a flat substrate from the same elastic material (Young's modulus $E_r$ and Poisson's ratio $\nu_r$). In our implementation this assembly is produced from more rigid material and forms the top part of the Elastomeric Pillar Cages (EPC). At its apex this micropillar firmly adheres to a flat elastic substrate (Young's modulus $E_s$, Poisson's ratio $\nu_s$). In EPCs this substrate is very soft, hence the subscript 's'. Moreover, we replace the distributed cell forces by a single point force $F_E$ (E for 'external') that acts at a distance $l_E$ from the surface of the rigid top substrate. For the geometry see Fig. 1.

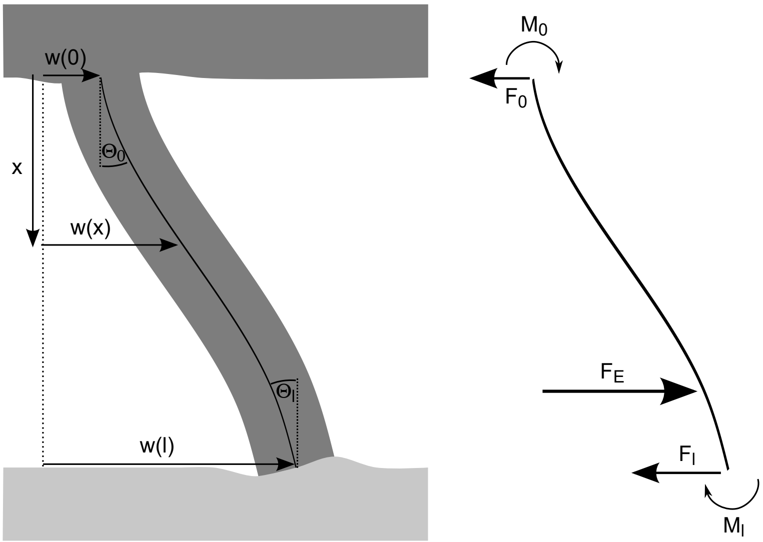


Figure 1: Left: Basic layout underlying our calculations. A micropillar (length $l$, radius $r$) emerges from an elastic substrate (Young's modulus $E_r$, Poisson's ratio $\nu_r$; dark grey). It firmly adheres to another elastic substrate (Young's modulus $E_s$, Poisson's ratio $\nu_s$, light grey). It is deformed by an external point force $F_E$ that acts at a distance $l_E$ from the rigid substrate. Distance from the rigid top is denoted by $x$, deflection of the neutral axis from the force free state (straight line, dotted), $w(x)$, is calculated. Right: Acting forces and moments are the external force $F_E$, the moment $F_E l_E$ connected to it, and, at both ends of the micropillar, counterforces $F_0$ and $F_l$ as well as bending moments $M_0$ and $M_l$.

Because the external force $F_E$ attacks at a distance from the substrate surfaces, a corresponding moment is applied to the beam. Force and moment are balanced by forces and moments at both ends of the micropillar. These arise from the deformations of the flat subtrates. In mechanical equilibrium forces and moments must balance.

$$F_0 + F_E + F_l = 0 \tag{1}$$

$$M_0 + F_E l_E + M_l + l F_l = 0 \tag{2}$$

For the balance of moments, Eq. 2, we omitted the force at $l = 0$ because it attacks at the origin of the coordinate system. One of the central results of Euler-Bernoulli theory of beam bending is that the deflection of the beam's neutral axis, $w(x)$, results from the bending moment, $M(x)$, acting in the beam at position $x$ according to the following differential equation

$$w'' = -\frac{M(x)}{E_r I}. \tag{3}$$

Here the apostroph signifies derivation with respect to $x$ and $I$ the beam's cross sectional moment of inertia. For a circular cross-section the latter is given by $\pi r^4/4$. The moments acting in this beam are given by

$$\begin{array}{lrcl} \text{For } x < l_E & M(x) & = & M_0 - xF_0 \\ \text{For } x > l_E & M(x) & = & M_0 - xF_0 - (x - l_E)F_E \end{array} \tag{4}$$

Given two boundary conditions (usually for positions and / or tangents at the endpoints) Eqs. 3 and 4 result in a unique solution for the shape of the neutral axis. Unfortunately, for this specific geometry these boundary conditions must be determined in an indirect way. Our strategy towards this end is as follows.

Because this mechanical system is statically indeterminate, the bearing reactions $F_0$, $F_l$, $M_0$, and $M_l$ must be determined explicitely. Here we use approximations for the distribution of forces in slender beams (Euler-Bernoulli beams) together with the well developed theory for the deformation of an elastic half space under the influence of surface forces, see [38,39]. From these approximations we find that the moment applied at a foundation of the beam is directly proportional to the tangent angle at this end whereas force is directly proportional to displacement. In other words, at both ends we find Hookean behavior with respect to torque and force. Therefore values and first derivatives of the solution of Eqs. 3 and 4 at both end points are equivalent to the acting forces and moments. We solve the differential equation for the neutral axis using force and momentum at one end as arbitrary parameters, which results in an explicit relation connecting $F_l$ and $M_l$ to $F_0$ and $M_0$. This is than inserted into force and momentum balance, Eqs. 1 and 2. This procedure results in two equations with two unknowns and, therefore, in a unique solution.

#### 3 Approximate Bearing Stiffnesses

The deformations of an elastic half space under the influence of a force distribution at its surface can be easily calculated by convolution of the force distribution with the displacement field caused by a point force [38,39]. Based on this Maloney et al. [40] calculated the displacements caused by a constant force that acts in a circular region of the interface. They give an explicit equation for the displacement of the center of the circle that we use here to estimate the stiffness of the beam's foundation against tangential forces.

$$\begin{array}{rclcrcl} F_0 & = & -k_0 w(0) & \text{with} & k_0 & = & \frac{\pi r E_r}{(1+\nu_r)(2-\nu_r)} \\ F_l & = & -k_l w(l) & \text{with} & k_l & = & \frac{\pi r E_s}{(1+\nu_s)(2-\nu_s)} \end{array} \tag{5}$$

Obviously, the beam connects to an elastic material of finite thickness. Some of us have given the displacement field for a point force acting on an elastic layer of finite thickness [21]. Based on this result Maloney et al. [40] incorporated the effects of finite thickness $h$. They find a stiffening due to the finite layer thickness that decreases with increasing layer thickness. At a layer thickness of 3.42 times the radius of the force application region this stiffening has decreased to a mere 10% effect. Because in EPC layer thickness amounts to more than ten micropillar radii this stiffening effect can be safely neglected.

Some of us gave the following approximate relation between the bending moment acting on the foundation of a micropillar emerging from an elastic layer and its angle to the normal [42].

$$\begin{aligned} M_0 &= -\mu_0 w'(0) \qquad \text{with} \quad \mu_0 = \frac{2\pi E_r I}{3r\alpha} \\ M_l &= -\mu_l w'(l) \qquad \text{with} \quad \mu_l = \frac{2\pi E_s I}{3r\alpha} \end{aligned} \tag{6}$$

Here $I$ denotes again the cross-sectional area of momentum of the beam and the numerical factor $\alpha$ amounts to 1.067. This equation was derived from the force distribution acting in a bent slender beam far away from the ends applied to the surface of an elastic half space. Here as well the convolution with the point force response was used. Obviously, the situation in the real bearing is more complicated. Therefore Kajzar et al. tested the validity of Eq. 6 on a macroscopic model and found indeed a proportional dependence of angle on torque. The experimental proportionality factor was about 7% smaller than $\alpha$ [42]. Compared to the uncertainties introduced by the measurement of the parameters (e.g. $r$, $E_r$, $E_s$) this level of approximation is well acceptable.

#### 4 The Neutral Axis

Equations 3 and 4 are easily integrated. At the point of force application, $l_E$, the solution $w$ and its first derivative $w'$ must be continuous. The solution is

$$w = \begin{cases} a_1 + a_0 x - \frac{1}{2}\frac{M_0}{E_r I}x^2 + \frac{1}{6}\frac{F_0}{E_r I}x^3 & \text{for} \quad x < l_E \\ a_1 + a_0 x - \frac{1}{2}\frac{M_0}{E_r I}x^2 + \frac{1}{6}\frac{F_0}{E_r I}x^3 + \frac{1}{6}\frac{F_E}{E_r I}(x - l_E)^3 & \text{for} \quad x > l_E \end{cases} \tag{7}$$

At $x = 0$ $w = a_1$ and $w' = a_0$. Therefore $a_1$ corresponds to the tangential force acting at the beam's foundation and $a_0$ to momentum.

#### 5 Determination of the Bearing Reactions

As stated above, we find the integration constants of the differential equation from force and momentum acting at $x = 0$.

$$\begin{aligned} a_0 &= -M_0/\mu_0 \\ a_1 &= -F_0/k_0 \end{aligned} \tag{8}$$

Using force balance, Eq. 1, and the stiffness at the softer interface, $k_l$, Eq. 5, we obtain

$$w(l) = (F_0 + F_E)/k_l \tag{9}$$

which yields together with Eq. 7 for the neutral axis and Eq. 8 a first linear equation that relates the bearing reactions $F_0$ and $M_0$.

$$\begin{aligned} F_0\left[1/k_0 + 1/k_l - l^3/(6IE_r)\right] + M_0 l\left[1/\mu_0 + l/(2IE_r)\right] + \\ + F_E\left[1/k_l - (l - l_E)^3/(6IE_r)\right] = 0 \end{aligned} \tag{10}$$

Now, we use similar procedures to the momentum balance. Using it, Eq. 2, the force balance, Eq. 1, and the stiffness against torque at the softer interface, $\mu_l$, see Eq. 6, we obtain

$$w'(l) = [M_0 - F_0 l - F_E (l - l_E)] / \mu_l \tag{11}$$

which yields together with Eq. 7 for the neutral axis and Eq. 8 a second linear equation that relates the bearing reactions $F_0$ and $M_0$.

$$\begin{aligned} -F_0 l [1/\mu_l + l/(2IE_r)] + M_0 [1/\mu_0 + 1/\mu_l + l/(IE_r)] & \\ -F_E (l - l_E) [1/\mu_l + (l - l_E)/(2IE_r)] &= 0 \end{aligned} \tag{12}$$

Together Eqs. 10 and 12 form a set of two linear equations for the two unkowns $F_0$ and $M_0$. These solutions exist but, unfortunately, they exhibit little obvious structure besides the fact that $F_0$ and $M_0$ are directly proportional to the external force $F_E$. As the solutions are best calculated using a computer algebra system, we refrain from quoting the very lengthy results. Instead we reproduce the Maple (Maple 2018.2, Maplesoft, Waterloo, Ontario, Canada) code we used for determining the solution of the equation set 10 and 12.

```
EI:=Er*MI;
A1:=1/k0 + 1/kl - 1/6*l^3/EI;
A2:=l*(1/m0 + 1/2*l/EI);
A3:=1/kl - 1/6*(l-lE)^3/EI);
B1:=-l/ml - 1/2*l^2/EI;
B2:=1/m0 + 1/ml + l/EI;
B3:=-(l-lE)/ml - 1/2*(l-lE)^2/EI;
solutions:=solve({A1*F0 + A2*M0 + A3*FE=0, B1*F0 + B2*M0 + B3*FE=0},[F0,M0]);
assign(solutions);
```

Into these solutions Eqs. 5, 6, and the moment of inertia are inserted and the neutral axis, Eq. 7, is calculated explicitely. As both, $F_0$ and $M_0$, are directly proportional to the external force the displacements are also.

#### 6 Structure of the Solution

In Fig. 2 we show the shape of the normalized neutral axis (deflection divided by $F_E$). Typical for the behavior of slender beams is the dependence on the position of force application, $l_E$. Here we observe a massive softening of the response with force application further away from the more rigid substrate. Intriguing and specific for a beam that is anchored at both ends is that the maximum deflection moves away from the apex of the beam if force is applied closer to the more rigid substrate, in the example of Fig. 2, left, at $l_E$ of 30 µm or less. Please note that this maximum is not occuring at the point of force application, instead it is shifted towards the softer substrate. Therefore a hypothetical cell that applies a force at one point will experience a compliance that is not determined by the maximum of the neutral axis, i.e. the maximum deflection, but by the deflection of the micropillar at the point of force application. The course of this

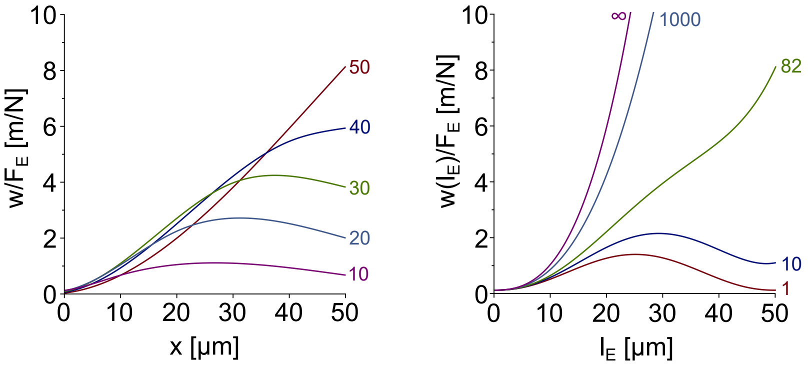


Figure 2: Behavior of the normalized deflection of the micropillar. Parameters throughout: $l = 50$ μm, $r = 5$ μm, $E_r = 1.3$ MPa, $\nu_r = \nu_s = 1/2$. Left: Neutral axes, $w(x)$, divided by acting force, $F_E$. $E_s = 15$ kPa, numbers at the individual curves give $l_E$ in μm. Right: The normalized deflection of the micropillar at the point of force application, $w(l_E)/F_E$, numbers at curves give $E_r/E_s$.

experienced or effective compliance is also shown in Fig. 2. If both flat substrates are of comparable stiffness, the experienced compliance displays a pronounced maximum at intermediate heights, whereas at very large differences between both stiffnesses, the experienced compliance is highest at the apex in contact with the soft substrate and approaches the behavior expected for a usual micropillar.

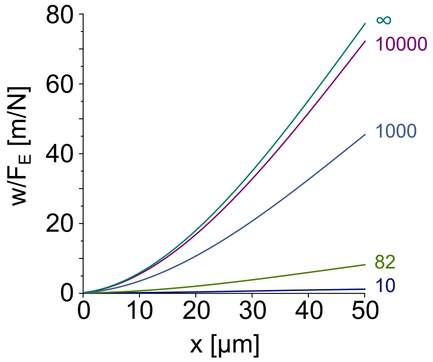


Figure 3: Normalized deflections of the micropillars (neutral axes, $w(x)$, divided by acting force, $F_E$). Parameters: $l = 50$ μm, $r = 5$ μm, $l_E = 50$ μm, $E_r = 1.3$ MPa, $\nu_r = \nu_s = 1/2$, numbers at curves give $E_r/E_s$.

With decreasing rigidity of the soft substrate displacements of the micropillar increase strongly. However, even at extrem softness, in the example of Fig. 3 at 130 Pa, displacements are still lower than for a micropillar that is not connected to a second substrate (in Figs. 2 and 3 indicated by '∞').

For a further comparison to the familiar behavior of a micropillar with one free end we note that in this case the full force ($F_E$) and the full moment ($F_E l_E$) are transmitted to the more rigid substrate, i.e. here $F_0 = -F_E$ and $M_0 = -F_E l_E$. These values were compared with the force, $F_0$, and moment, $M_0$, transmitted to the more rigid substrate. In this procedure the stiffening effect of the soft substrate became very obvious. At the values realized in EPC ($E_r/E_s = 82$) all the 'soft' part of the system is dominated by force and momentum transfer to the soft substrate. Only at extremely low Young's moduli of the softer substrate the system approaches the behavior of a micropillar with one free end.

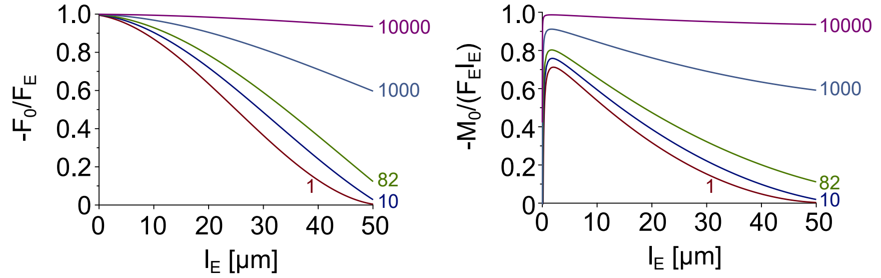


Figure 4: Force and momentum transfer to the rigid substrate. Parameters throughout: $l = 50$ µm, $r = 5$ µm, $E_r = 1.3$ MPa, $\nu_r = \nu_s = 1/2$. Numbers at curves give $E_r/E_s$. Left: force transfer. Right: momentum transfer. For a micropillar with one free end relative force and momentum transfer amount to exactly 1.

#### 7 Typical Values and Error Budget

For practical purposes the most important result of our calculation is that external force, $F_E$, and displacement, $w(x)$, are directly propotional, albeit with a lengthy parameter of proportionality that depends on many parameters. With the above equations it is straightforward to calculate this proportionality constant and, thus, forces from displacements. Estimates for the uncertainties of measured forces, $\Delta F$, were calculated using Gaussian error propagation.

$$\frac{\Delta F}{F} = \sqrt{\sum_{i=1}^{9} b_i^2} \quad \text{with} \quad b_i = \frac{dF}{da_i}\frac{\Delta a_i}{F} \tag{13}$$

Where $a_i$ denote the nine different parameters and $\Delta a_i$ their uncertainties.

In Table 1 we present typical parameter values and their uncertainties. For $r$, $E_r$, and $E_s$ averages and standard deviations from repeated calibrations were used. For $l$ the average micropillar height from repeated preparations was used, uncertainty is given by the accuracy of focussing the apex of the microcolumn. Poisson ratios were determined by stretching of cylindrical test pieces [41]. We found a value of 0.5, as expected for ultrasoft elastomers. The uncertainty of 0.02 is an estimate. The measurement height, $x$ was set to be 45 µm, its uncertainty is determined by how exact the surface of the flat substrate part can be focused. In this example we arbitrarily assumed a deflection of 1 µm, its uncertainty was estimated from the accuracy of relocalization of the fluorescent marker beads in the microcolumns (see main text). A major source of uncertainty is the height of force application, $l_E$ because neither cell adhesion complexes nor the force

| Parameter | Mean Value | Uncertainty | $\|\|b_i\|\|$ |
|---|---|---|---|
| $r$ | 5.03 µm | 0.25 µm | 7.77% |
| $l$ | 51.4 µm | 2 µm | 3.70% |
| $l_E$ | 45 µm | 5 µm | 8.41% |
| $E_r$ | 1.27 MPa | 0.16 MPa | 2.49% |
| $E_s$ | 15.5 kPa | 2.2 kPa | 11.4% |
| $x$ | 45 µm | 0.5 µm | 0.84% |
| $w$ | 1 µm | 0.1 µm | 10% |
| $\nu_r$ | 0.5 | 0.02 | $4 \cdot 10^{-15}$ |
| $\nu_s$ | 0.5 | 0.02 | $6 \cdot 10^{-22}$ |

Table 1: Average values of parameters, their uncertainties, and contributions to the overall measurement uncertainty. For definition of $b_i$, see Eq. 13. At these specific parameters, deflection $w$ of 1 µm corresponds to a force of 162 nN. Overall uncertainty is in this example 19.5%.

producing cytoskeleton could be imaged. Our estimate simply arises from the maximum height of a cell that is adhered to the soft substrate, a correspondingly large uncertainty was assumed.

The overall uncertainty of force measurement of 19.5% is dominated by the contributions of $E_s$ (11.4%), $w$ (10%), $l_E$ (8.4%), and $r$ (7.8%). The large uncertainty contributed by the elasticity of the soft substrate reflects the experimental challenges in the preparation of ultrasoft elastomers. The overall uncertainty is similar to the one given by Kajzar et al. for force measurement with free micropillars [42]. In view of these significant experimental uncertainties a more elaborate treatment of the mechanics of this system, e.g. by finite element simulations or by including rounding of the beam's apex, seems not justified.

Calculated from the parameters given in Table 1 the stiffness of EPC for a force applied at the soft substrate, i.e. the point of lowest stiffness, is 162 mN/m, well in the range of soft cantilevers for surface force microscopy.

#### Bibliography

The following references from the main text are cited in this Appendix.

[21] R. Merkel, N. Kirchgeßner, C. M. Cesa, and B. Hoffmann. Cell force microscopy on elastic layers of finite thickness. *Biophysical Journal*, 93:3314–3323, 2007.

[38] J. Boussinesq. *Application des potentiels à l'étude de l'équilibre et du mouvement des solides élastiques*. Gauthier-Villars, Paris, F, 1885.

[39] L. D. Landau and E. M. Lifshitz. *Course of Theoretical Physics. VII Theory of Elasticity*. Pergamon Press, Oxford, UK, 1986.

[40] J. M. Maloney, E. B. Walton, C. M. Bruce, and K. J. Van Vliet. Influence of finite thickness and stiffness on cellular adhesion-induced deformation of compliant substrata. *Physical Review E*, 78:041923, 2008.

[41] C. M. Cesa, N. Kirchgeßner, D. Mayer, U. S. Schwarz, B. Hoffmann, and R. Merkel. Micropatterned silicon elastomer for high resolution analysis of cell force patterns. *Reviews of Scientific Instruments*, 78:034301, 2007.

[42] A. Kajzar, C. M. Cesa, N. Kirchgeßner, B. Hoffmann, and R. Merkel. Towards physiological conditions for cell analyses: Forces of heart muscle cells suspended between elastic micropillars. *Biophysical Journal*, 94:1854–1866, 2008.

[43] S. P. Timoshenko and J. N. Goodier. *Theory of Elasticity*. McGraw-Hill, Auckland, NZ, 1970.
