## Supplementary Information for "Elastomeric Pillar Cages Modulate Actomyosin Contractility of Epithelial Microtissues by Substrate Stiffness and Topography"

**Movie S1:** Lateral cell injection of MCF10A cells using a glass microcapillary.

**Movie S2:** Laser ablation (white arrowhead) of a myofibroblast (labeled by MitoTracker dye). Pillars with incorporated QDots (gray) were used for displacement tracking.

**Movie S3:** Time lapse of spontaneous beating cardiomyocytes (labeled by MitoTracker dye, orange). Pillars with incorporated QDots (green) were used for displacement tracking (Imaging interval: 90 ms).

**Appendix A1**: Analytical approximation of EPC-derived cell forces.

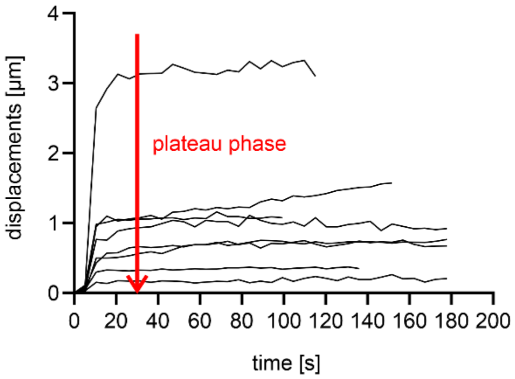


**Figure S1.** Time-dependent pillar relaxation. Shown is the displacement of single pillars over time after MFB cell ablation. Displayed are the displacements of single pillars caused by single cells that adhered to single pillars (n = 8). QDot displacements were measured at 5 µm pillar height (above the bottom layer). The red arrow indicates the beginning of the plateau phase at t = 30 s after ablation. This time point was chosen for all force measurements to minimize the influence of unspecific pillar drift.

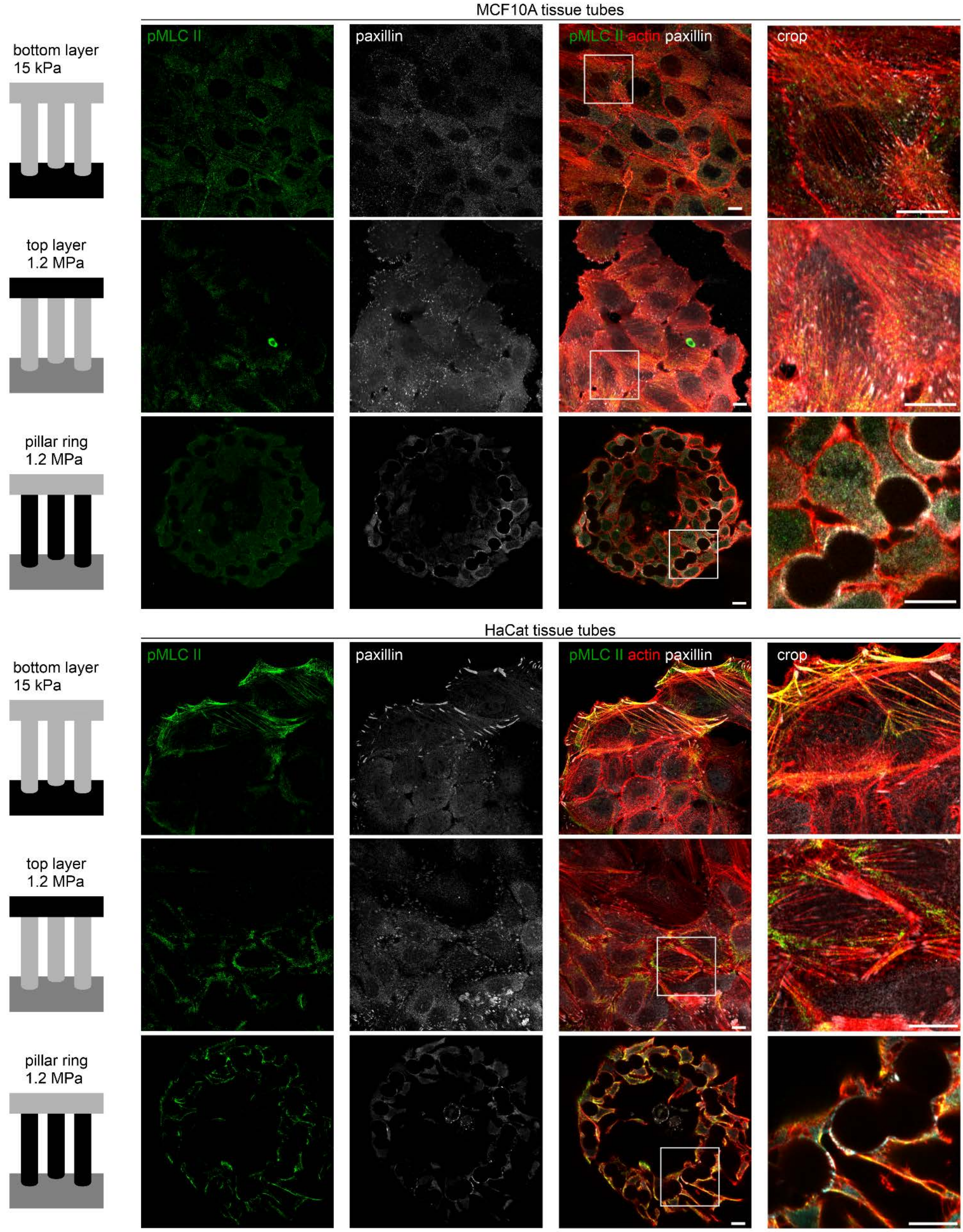


**Figure S2**. Modulation of the actomyosin cytoskeleton by the EPC geometry. Micrographs show the distinct cellular localization of actin stress fibers with phosphorylated myosin light chain II (pMLC II) in MCF10A and HaCat microtissues, depending on cell location within the EPC topography. Microtissues were grown for seven days, fixed and stained against pMLC II (green), focal adhesion marker paxillin (gray) and actin cytoskeleton (phalloidin, red). Imaging plane: 10 µm pillar height. Scale bars = 10 µm.
